# Sex chromosome differentiation via changes in the Y chromosome repeat landscape in African annual killifishes *Nothobranchius furzeri* and *N. kadleci*

**DOI:** 10.1101/2022.03.13.484186

**Authors:** Jana Štundlová, Monika Kreklová, Karolína Lukšíková, Anna Voleníková, Tomáš Pavlica, Marie Altmanová, Martin Reichard, Martina Dalíková, Šárka Pelikánová, Anatolie Marta, Sergey A. Simanovsky, Matyáš Hiřman, Marek Jankásek, Tomáš Dvořák, Joerg Bohlen, Petr Ráb, Petr Nguyen, Alexandr Sember

**Affiliations:** Laboratory of Fish Genetics, Institute of Animal Physiology and Genetics, Czech Academy of Sciences, Liběchov, Czech Republic; University of South Bohemia, Faculty of Science, České Budějovice, Czech Republic; Department of Genetics and Microbiology, Faculty of Science, Charles University, Prague, Czech Republic; Department of Zoology, Faculty of Science, Charles University, Prague, Czech Republic; Department of Ecology, Faculty of Science, Charles University, Prague, Czech Republic; Institute of Vertebrate Biology, Czech Academy of Sciences, Czech Republic; Department of Ecology and Vertebrate Zoology, University of Łódź, Łódź, Poland; Department of Botany and Zoology, Masaryk University, Brno, Czech Republic; Biology Centre of the Czech Academy of Sciences, Institute of Entomology, České Budějovice, Czech Republic; Institute of Zoology, Chisinau, Moldova; Severtsov Institute of Ecology and Evolution, Russian Academy of Sciences, Moscow, Russia

**Keywords:** interpopulation variability, inversion, recombination suppression, sex chromosome degeneration, repeatome, RepeatExplorer, sex chromosome polymorphism, synaptonemal complex, Teleostei

## Abstract

Repetitive DNA represents an important driver of sex chromosome differentiation. Yet, repetitive sequences tend to be misrepresented or overlooked in genomic studies. We analysed repetitive DNA landscape of sex chromosomes in several populations of a turquoise killifish *Nothobranchius furzeri* and its sister species *N. kadleci* (Teleostei: Nothobranchiidae), representatives of African annual killifishes with high rate of karyotype and sex chromosome evolution. We combined bioinformatic analyses of repeatome with molecular cytogenetic techniques such as comparative genomic hybridization, fluorescence *in situ* hybridization with satellite sequences, genes for ribosomal RNAs (rDNA) and bacterial artificial chromosomes (BACs) and immunostaining of SYCP3 and MLH1 proteins, which marked lateral elements of synaptonemal complexes and recombination sites, respectively. We revealed that *N. furzeri* and *N. kadleci* share the XY sex chromosome system, which is thus much older than previously assumed. Sex chromosomes are mostly heteromorphic as evidenced by distinct distribution of satellite DNAs and major rDNA. Yet, the heteromorphic XY sex chromosomes pair almost exclusively regularly in meiosis, which implies synaptic adjustment. Physical mapping of BACs identified inversions on Y chromosomes of the *N. kadleci* populations, akin to the pattern previously reported in *N. furzeri*. Yet, repetitive DNA landscape of X and Y sex chromosomes either diverged in parallel in populations of both species or it evolved in their common ancestor and thus predates the inversions. The observed differentiation via repeat repatterning thus cannot be explained by the classical sexually antagonistic model. Rather, we hypothesized that relaxed meiotic drive and recombination reduced by neutral processes could drive changes in repeatome and secondary inversions could be maintained by sexually antagonistic regulatory effects resulting from early evolution of dosage compensation.

**Author summary:** Early differentiation of sex chromosomes is not yet satisfactorily understood despite intensive research effort. Homomorphic sex chromosomes and their rapid turnover are common in teleost fishes, which makes them excellent models for studying evolution of nascent sex chromosomes. We investigated sex chromosomes in several populations of two sister species of African annual killifishes, *Nothobranchius furzeri* and *N. kadleci*, particularly their repetitive landscape, which was misrepresented in previous genomic studies. Combination of cytogenetic and genomic approaches revealed that both species share heteromorphic XY sex chromosome system. The *N. furzeri* XY sex chromosomes thus evolved earlier than previously expected. In *N. kadleci*, Y-linked inversions analogous to those reported in *N. furzeri* were detected. Changes in repetitive DNA distribution on sex chromosomes are either convergent or occurred in a common ancestor of both species, prior to the inversion events. The observed sex chromosome differentiation on repetitive DNA level thus cannot be reconciled with the classical theoretical model of sex chromosome evolution driven by sexually antagonistic selection. We invoke alternative explanations such as relaxed meiotic drive and recombination reduced by neutral processes, and we hypothesize that secondary inversions could be maintained by early evolution of dosage compensation resulting in sexually antagonistic regulatory effects.

## Introduction

Sex chromosomes are among the most dynamic parts of genomes. They emerged repeatedly and independently in diverse eukaryotic lineages, yet their evolution shares common features [1–4]. These specialized chromosomes usually evolve from an autosomal pair after one homologue acquires a master sex determining (MSD) gene [5–7]. The classical theoretical model assumes recombination is reduced in the newly established sex-determining (SD) region due to sexually antagonistic selection, which favours a tight genetic linkage of the MSD gene with sexually antagonistic alleles, i.e. allele beneficial to one sex but detrimental for the other. However, empirical evidence supporting the sexual antagonism (SA) hypothesis is scarce and alternative, mostly neutral models have recently been proposed [3,8–11]. In any case, the non- recombining region on the sex-limited chromosome, i.e. Y in male heterogametic or W in female heterogametic systems, starts accumulating mutations and repetitive sequences. The non-recombining region can further spread due to either genic modifiers or chromosome rearrangements mediated by repeats. Deletions, duplications, and insertions of repetitive sequences may also affect the size of sex-limited chromosomes. Altogether, these processes can eventually produce a heteromorphic sex chromosome pair, which is detectable cytogenetically [4,6,13–15].

In teleost fishes, about 500 sex chromosome systems have been identified so far 16–19] and references therein). Regardless of their evolutionary age, the majority of these systems is homomorphic, i.e. they show low levels of differentiation, and hence are indistinguishable by standard cytogenetic protocols [14,16,17]. Teleost sex chromosomes usually differ considerably between closely related species and even between conspecific populations. The high rate of sex chromosome turnover [14,16,18,20–22] makes fish sex chromosomes an excellent model system for investigation of early phases of sex chromosome differentiation, drivers of the sex chromosome turnover and the role of these processes in speciation and adaptation of fishes [16,23–27].

African killifishes of the genus *Nothobranchius* Peters, 1868 (Aplocheiloidei: Nothobranchiidae) are small-bodied freshwater fishes with pronounced sexual dimorphism and dichromatism (males are larger and more colourful) [28]. The genus *Nothobranchius* currently contains more than 90 species [29], which are adapted to periodically dry pools in the south- east African savannahs. Due to the temporarity of this environment, *Nothobranchius* (as well as five other clades within the order Cyprinodontiformes) evolved eggs that undergo diapause to endure the dry period [30]. The wet season in these pools lasts only few months, enforcing the fish to hatch, grow, mature, and reproduce in a very short time [30, 31]. In the turquoise killifish *N. furzeri* Jubb, 1971, the wet part of the life cycle can be as short as 30–40 days [32]. This life cycle part is rather compressed than prematurely terminated, and the fish experience typical age-related vertebrate pathologies, which makes them a useful model to study the biology of aging [31, 33]. Most biomedical and genomic research has been addressed using the model species *N. furzeri*, where the transcriptomic and genomic resources as well as genome editing tools have been established [34–39]. Its sister species *N. kadleci* Reichard, 2010 [40, 41] received considerably less attention [42, 43].

*Nothobranchius* killifishes underwent a high number of karyotype changes, leading to a wide range of diploid chromosome numbers (2n = 16 – 50) and striking variability in karyotype composition found in 65 species examined to date [44]. It was shown that genomes of *Nothobranchius* spp. contain high amounts of repetitive DNA [35,45,46] which may facilitate karyotype evolution. Intriguingly, six *Nothobranchius* species possess X1X2Y multiple sex chromosome system [44,47,48]. Multiple sex chromosomes can be easily identified by means of cytogenetics, unlike the XY sex chromosome system from which they evolved. Indeed, the XY sex chromosome pair has only been confirmed in *N. furzeri* using a genomic survey [35]. Notably, the SD region varied in size between different *N. furzeri* populations [35, 39], ranging from 196 kb to 37 Mb [35]. It was proposed that the *N. furzeri* Y chromosome polymorphism represents an early stage of sex chromosome evolution. An Y-linked allele of *gdf6* gene (*gdf6Y*) has been identified as a putative MSD gene in the turquoise killifish [35].

In the present study, we investigated the origin of and processes shaping the sex chromosome differentiation in *N. furzeri.* Hence, we performed detailed cytogenomic analyses including comparative genomic hybridization (CGH) and physical mapping of various repeats and bacterial artificial chromosomes (BAC) in three populations of *N. furzeri* and two populations of its sister species, *N. kadleci* (Fig 1). Our results revealed that *N. furzeri* and *N. kadleci* share a heteromorphic XY sex chromosome system with the same putative MSD gene and highly similar patterns of differentiation, implying that this system has longer evolutionary history than previously anticipated. Y-linked inversion polymorphism described in *N. furzeri* [35] occurs also in *N. kadleci* but these events do not represent primary key drivers of XY sex chromosome evolution. We show that sex chromosome differentiation occurred via changes in the repetitive DNA landscape, which are independent from the evolution of the non-recombining SD region.

**Fig 1.**
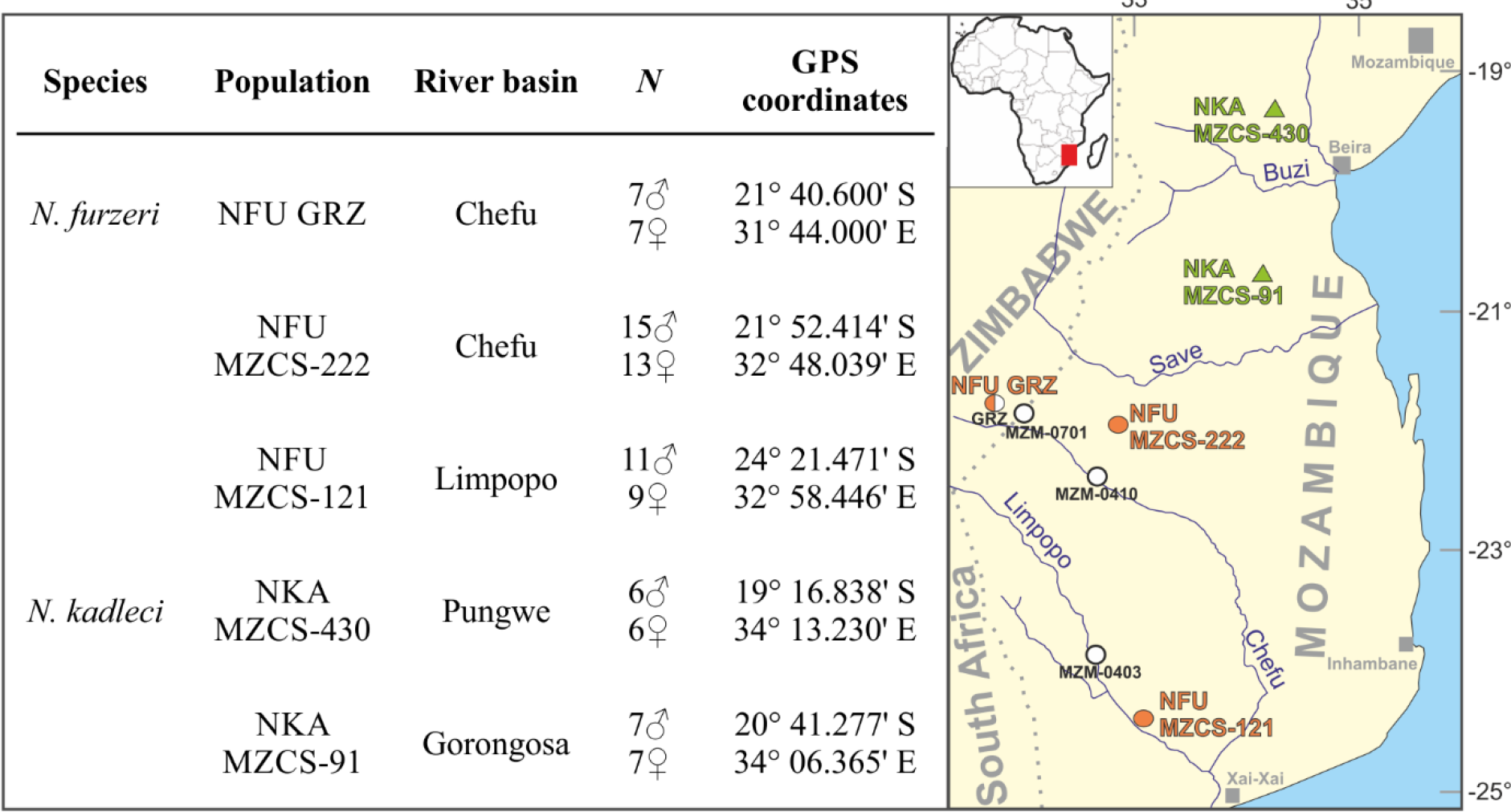
List of *N. furzeri* and *N. kadleci* killifish populations used in this study, with assignment to their phylogeographic lineage and sample sizes. The studied individuals (left panel) were samples from wild-derived captive populations (for details, see [43,49–51]. Right panel: Map with geographic origin of sampled populations of *N. furzeri* (orange circles) and *N. kadleci* (green triangles) as well as *N. furzeri* populations studied previously by Reichwald et al. [35], namely MZM-0701, MZM-0410, and GRZ of Chefu clade, and MZM-0403 of Limpopo clade (white circles). GRZ strain was researched both in Reichwald et al. [35] and in the present study (mixed white-orange circle). Abbreviations: *N* = number of analysed individuals.

## Results

### Basic karyotype characteristics

Karyotype of individuals from all studied *N. furzeri* and *N. kadleci* populations consisted of 19 chromosome pairs (S1 and S2 Figs) as reported previously [35,44,45]. Besides three metacentric, six submetacentric and nine subtelo-to-acrocentric (st-a) pairs, the female karyotype also contained one pair of metacentric (or submetacentric; NFU GRZ) X sex chromosomes and the male karyotype harboured a single X chromosome and one metacentric, submetacentric or subtelocentric Y chromosome. The presence of sex chromosomes, however, could not be revealed by conventional karyotyping alone. We inferred this information primarily from BAC-FISH experiments (see below).

Precise identification of chromosome pairs was challenging due to variable size and degree of spiralization of remarkably large blocks of (peri)centromeric heterochromatin. They were revealed by C-banding in all chromosomes regardless the sex, population and species (S3 Fig). Additional less prominent interstitial and terminal heterochromatin blocks were observed on some chromosomes including the Y sex chromosome. (Peri)centromeric heterochromatin blocks were further strongly stained by GC-specific fluorochrome Chromomycin A3 (S4 Fig).

### Characterization of repeat landscape

We employed RepeatExplorer2 pipeline [52] to analyse the repeatome of *N. furzeri* (GRZ) and *N. kadleci* (MZCS-430) genomes. In total, approximately 76 % of *N. furzeri* and 73 % of *N. kadleci* reads were clustered as repetitive DNA. We are, however, aware that RepeatExplorer2 often excludes microsatellites [52], therefore the obtained values highly likely do not represent the exact proportion of repetitive DNA in the genomes. The most abundant repeats (with estimated genome proportion at least 0.5 %) are listed in Table 1 and include both satellites and mobile elements. A major part of the repetitive DNA was composed of only two satellites, namely Nfu-CL1 and Nfu-CL2, covering ca. 19.5 % and 17.1 % of the genome of *N. furzeri* and *N. kadleci*, respectively. The abundance of the remaining repeats did not exceed 1%. For all repeats in Table 1, the corresponding counterparts were identified in both analysed genomes, suggesting that the repeatome is largely shared by *N. furzeri* and *N. kadleci* species, but differs in the abundance of the repetitive DNAs. In some cases, such as Nfu-CL4 and Nka-CL8, BLAST/N search yielded more than one significant hit. Detailed examination of the first one revealed that the Nfu-CL4 LINE retrotransposon was divided into several clusters in *N. kadleci* genome, however, all belonged to the same supercluster and thus most likely represented the same element. Interestingly, this was not the case for Nka-CL8 LTR retrotransposon, where contigs showed high similarity with three small independent *N. furzeri* clusters. This could be caused either by low amount of reads in these clusters, not sufficient to reconstruct the whole element, or its different organization in *N. furzeri* genome.

**Table 1.**
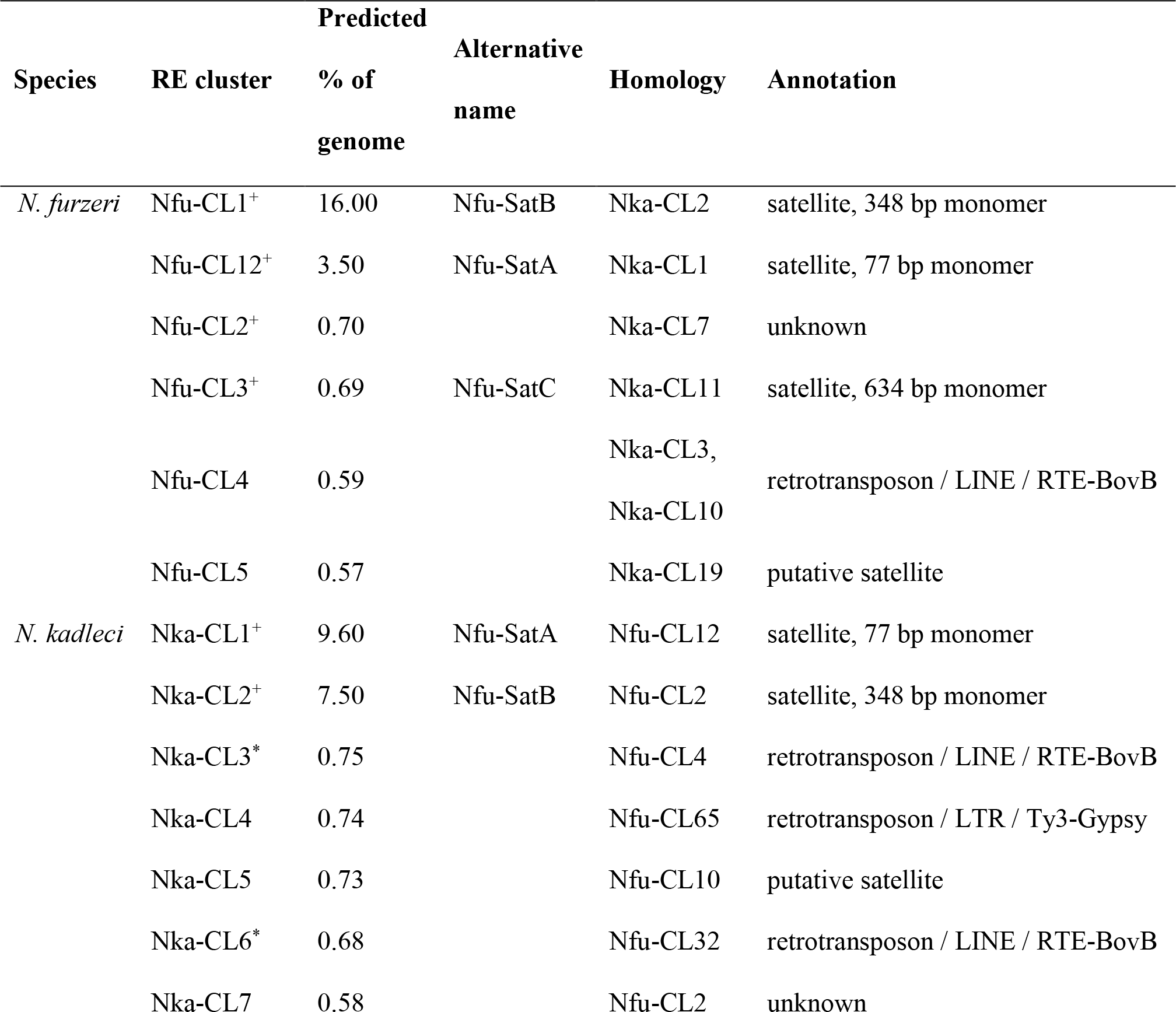
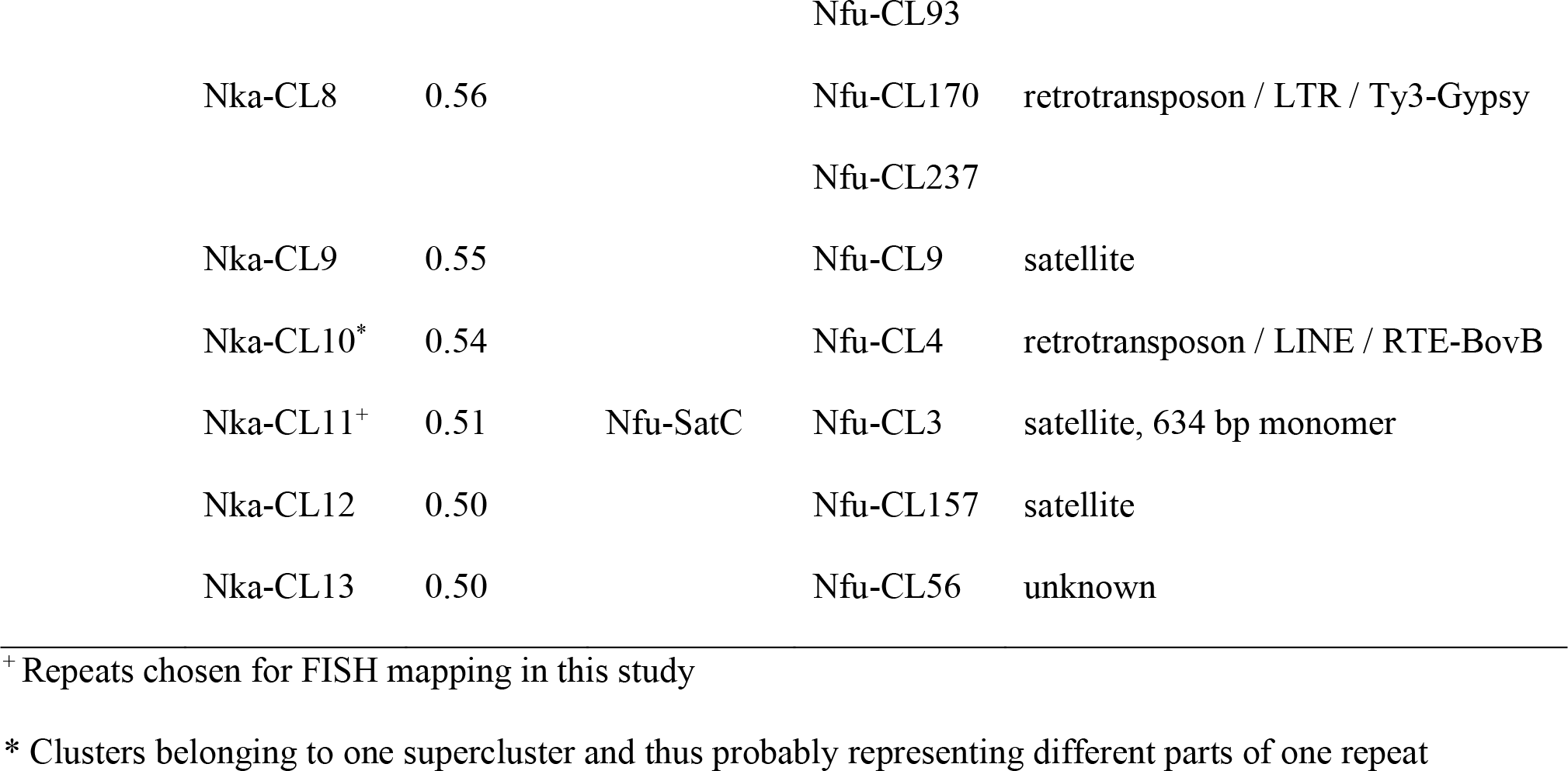
The most abundant repetitive sequences identified by RepeatExplorer2 pipeline in *Nothobranchius furzeri* and *N. kadleci* killifishes (≥0.5% genome)

### Major satellite markers

The tree satellites with the largest monomeric units in the *N. furzeri* genome, designated as Nfu-SatA (Nfu-CL12), Nfu-SatB (Nfu-CL1) and Nfu-SatC (Nfu-CL3), were chosen for chromosomal mapping and are described in further details in S1 Text. Briefly, Nfu-SatA and Nfu-SatB are the most abundant repeats in *N. furzeri* and *N. kadleci* genomes and were already previously identified and characterized in certain *N. furzeri* strains [35, 45]. Nfu-SatC was, given the characteristic size 634 bp, previously found to be enriched in Y-linked region of differentiation in *N. furzeri* [35].

### Hybridization patterns of repetitive DNA probes derived from RepeatExplorer2 analysis

Hybridization probes for Nfu-SatA and Nfu-SatB revealed virtually the same patterns across individuals and populations: large signals of different sizes in (peri)centromeric regions of all chromosomes regardless sex, population and species (S5 Fig). Only in the case of Nfu-SatB, male and female individuals from population NFU MZCS-121 displayed a notable gap in the signal in a single metacentric chromosome pair (S5H Figs). Similar gap was also observed after Nfu-SatC mapping (S6M Fig). In contrary with Nfu-SatA and Nfu-SatB, FISH with Nfu- SatC revealed a distribution in small clusters dispersed throughout the entire chromosome complement but with few more prominent accumulations (S5K–O Fig). One of these accumulations was especially remarkable and it was restricted to a single chromosome in males only, consistently in all analysed populations except for NFU MZCS-121.

### Sequence analysis of rDNA fragments

PCR amplification of 5S rDNA from the *N. kadleci* genomic DNA (gDNA) resulted consistently in ca. 500 bp long fragment. Sequencing of the cloned fragments followed by BLAST/N search at NCBI revealed that 470 bp long stretch of analysed sequence contains 94 nt of conserved fish 5S rDNA coding region (5’-truncated; the complete coding region has a conserved size 120 bp; [53]). The remaining sequence represents non-transcribed spacer (NTS) as confirmed by 96% sequence similarity of the entire 470 bp long fragment with the 5S rDNA sequences from *N. furzeri* genome (e.g sequence ID: EU780558.1.). Regarding 18S rDNA, for a majority of experiments we used optimized 18S rDNA probe previously prepared from *Hypophthalmichthys molitrix* [54]. For a subset of experiments, we prepared 18S rDNA probe from genomic DNA (gDNA) of *Nothobranchius guentheri*. We obtained 1200 bp long fragment which, when subjected to BLAST/N, displayed high similarity results (96-98% identity) with 18S rDNA sequences of many teleosts including those from *H. molitrix* (sequence ID: MT165584) and *N. furzeri* (sequence ID: EU780557.1). rDNA sequences were deposited in GenBank (accession numbers: MZ694985 for 5S rDNA from *N. kadleci* and MZ682111 for 18S rDNA from *N. guentheri*).

### Telomere probe mapping

FISH with PNA (peptide nucleic acid) telomere probe revealed signals at chromosomal ends invariably in all tested individuals across populations and regardless of sex. No deviations from this standard pattern were observed (S6 Fig).

### BAC-FISH and sequential analyses with a special focus on XY sex chromosome differentiation

In order to identify XY sex chromosomes we mapped four selected XY-linked BAC clones (S7 Fig) by FISH. All clones mapped consistently into the long (q) arms of a single chromosome pair (Figs 2 and S7). In males from most of analysed populations, the XY pair was formed by medium-sized metacentric X and a smaller subtelocentric Y element. In NFU GRZ population, X chromosomes were submetacentic. Y chromosome was submetacentric in NKA MZCS-430 population. In males from NFU MZCS-121 population, Y chromosome displayed variable morphology ranging from metacentric to submetacentric category and it was only slightly smaller than X chromosome counterpart (Figs 4, S7 and S8). In all females the pair bearing BAC-FISH signals was formed by metacentric (or submetacentric; NFU GRZ) Xs of the same or almost the same size.

**Fig 2.**
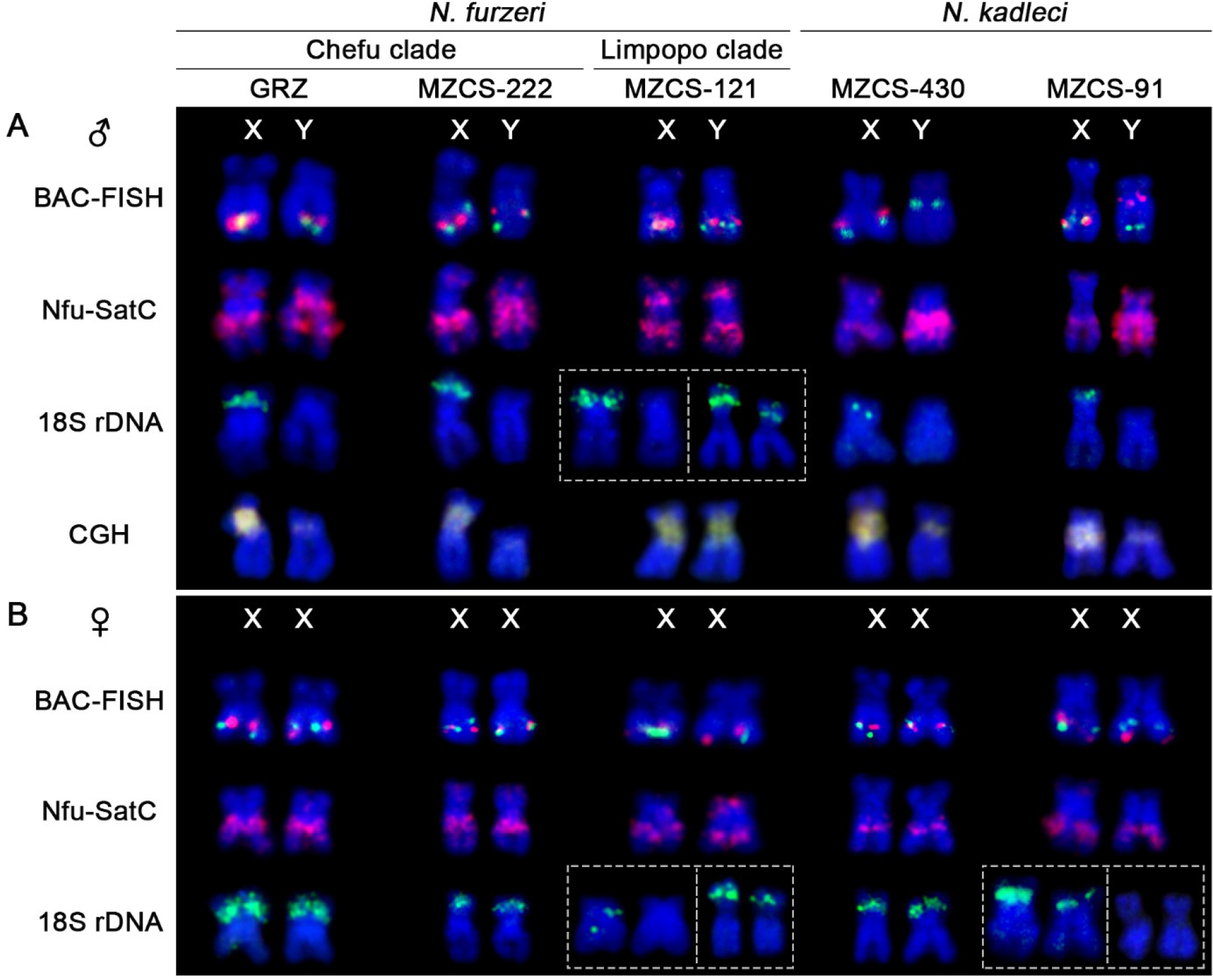
Partial karyotypes of XY and XX sex chromosomes from all *N. furzeri* and *N. kadleci* populations under study after sequential cytogenetic analyses. (A) males; (B) females. The same metaphase plates were hybridized subsequently with three rounds of FISH in the following order: i) BAC-FISH: dual-colour FISH with X-linked 220Bc03 (green signal) and 201Bd03 bearing *gdf6* gene (red signal). Note the shifted orientation of BACs on Y chromosome in both *N. kadleci* populations; ii) single-colour FISH with Nfu-SatC probe (red signals) – note contrasting accumulation of this repeat between X and Y sex chromosome (more pronounced in *N. kadleci* MZCS-430 and MZCS-91 and almost lacking in *N. furzeri* population MZCS-121); iii) 18S rDNA FISH (green signals). Note that except for NFU MZCS-121, 18S rDNA cluster is present on short arms of X chromosome but not on Y chromosome. Some *N. furzeri* males from Limpopo population MZCS-121 have exceptionally 18S rDNA site also on Y chromosome and some females from this population lacked 18S rDNA site on one X homologue. The polymorphic patterns are boxed. One or both 18S rDNA sites were absent on X chromosomes in some *N. kadleci* females from population MZCS-91 (full range of polymorphisms is present in S12 FigD); iv) comparative genomic hybridization (CGH) – hybridization patterns of male (red) and female (green) genomic probes yielded uniform yellowish signal. Note differences between X and Y in the length of (peri)centromeric block of repetitive DNA. Full karyotypes and hybridization patterns are provided in S8-S10 Figs.

In *N. furzeri* populations, all BAC clones showed identical location on both homologues regardless their size differentiation. In both populations of *N. kadleci*, however, the position of three BAC clones, specifically 262Ad08, 225De03 and 201Bd03 (*gdf6*-bearing clone), was remarkably shifted close to the centromere on Y but not on X chromosome. A clone 220Bc03 displayed also this Y-linked pattern but in NKA MZCS-430 population only, while in NKA MZCS-91 population it exhibited the same location as on X chromosome (S7 Fig).

rDNA FISH on reprobed metaphases of *N. furzeri* and *N. kadleci* individuals showed X-linked 18S rDNA locus located subterminally on the short (p) arms. This region lacked on Y chromosome (Figs 2, S8, S9 and S10). Nonetheless, in some males from NFU MZCS-121 population, 18S rDNA probe revealed signal also on equivalent location on Y chromosome, with inter-individual size differences (S8 Fig). This type of polymorphism was apparent also on preparations treated solely by rDNA FISH (S11E Fig). Finally, some females from NFU MZCS-121 and NKA MZCS-91 lacked 18S rDNA cluster on one X homologue and in a rare case of one NKA MZCS-91 female, both Xs lacked this region (S9-S12 Figs).

### Chromosomal distribution of 5S and 18S rDNA clusters

5S rDNA probe mapped consistently to the interstitial region of a single large submetacentric chromosome (pair No. 4 in *N. furzeri* and No. 3 in *N. kadleci*) in both sexes and no deviations from this pattern were revealed across populations of both species (S8-S12 Figs). On the other hand, mapping of 18S rDNA probe showed considerable polymorphisms in terms of cluster size and its presence/absence between homologs across different populations (S8-S12 Figs). Detailed description of these patterns is given in S2 Text.

### Molecular divergence of Y chromosomes as revealed by CGH

We applied CGH method in male-to-female comparison arrangement in each population as this approach might reveal sex chromosomes at certain stage of differentiation if the accumulated repeats are divergent enough qualitatively and/or quantitatively between sexes ([55, 56] and references therein). In our case, there was no differential pattern that could be revealed on sex chromosomes. However, CGH marked Y sex chromosome in the way that it was the only element in the complement which lacked huge (peri)centromeric accumulation in all but NFU MZCS-121 population (S8 and S10 Figs).

### Immunostaining and FISH analysis of synaptonemal complexes in male and female meiosis

We performed immunodetection of synaptonemal complex protein SYCP3 (lateral elements of synaptonemal complexes) on male pachytene spreads in order to analyse the chromosome pairing to seek for abnormalities of synapsis which may indicate the presence of region with suppressed recombination (and hence sex chromosomes). In males of all species, we observed 19 fully synapsed standard bivalents (Figs 3, S13 and S14). No consistent signs of delayed pairing, asynapses or other irregularities were observed (a rare exception can be seen in S13B Fig). Crossover sites were identified by immunolocalization of mismatch repair protein MLH1. We observed usually one or two (rarely three) MLH1 foci per bivalent. In majority of bivalents (especially those with two MLH1 foci) they were located close to chromosomal termini. Single interstitial MLH1 site per bivalent was present in a few bivalents only (Figs 3 and S13). To compare male and female MLH1 patterns, we investigated females from NFU MZCS-222 population and found highly similar patterns, with varied ratio of bivalents with one or more MLH1 foci and generally only slightly higher proportion of bivalents with a single MLH1 signal in the interstitial position compared to males (Fig 3C). Moreover, we observed unusual small bivalent-like structure without MLH1 signal in all studied females (Fig 3C). Very rarely, we observed such element in few male plates (data not shown). As our observation is restricted only to several females from *N. furzeri* population MZCS-222, we cannot make any conclusive assessment what this element might represent but it is strikingly reminiscent of supernumerary (B) and/or germline-restricted chromosomes (e.g. [57, 58]). This feature will require deeper investigation in future studies.

**Fig 3.**
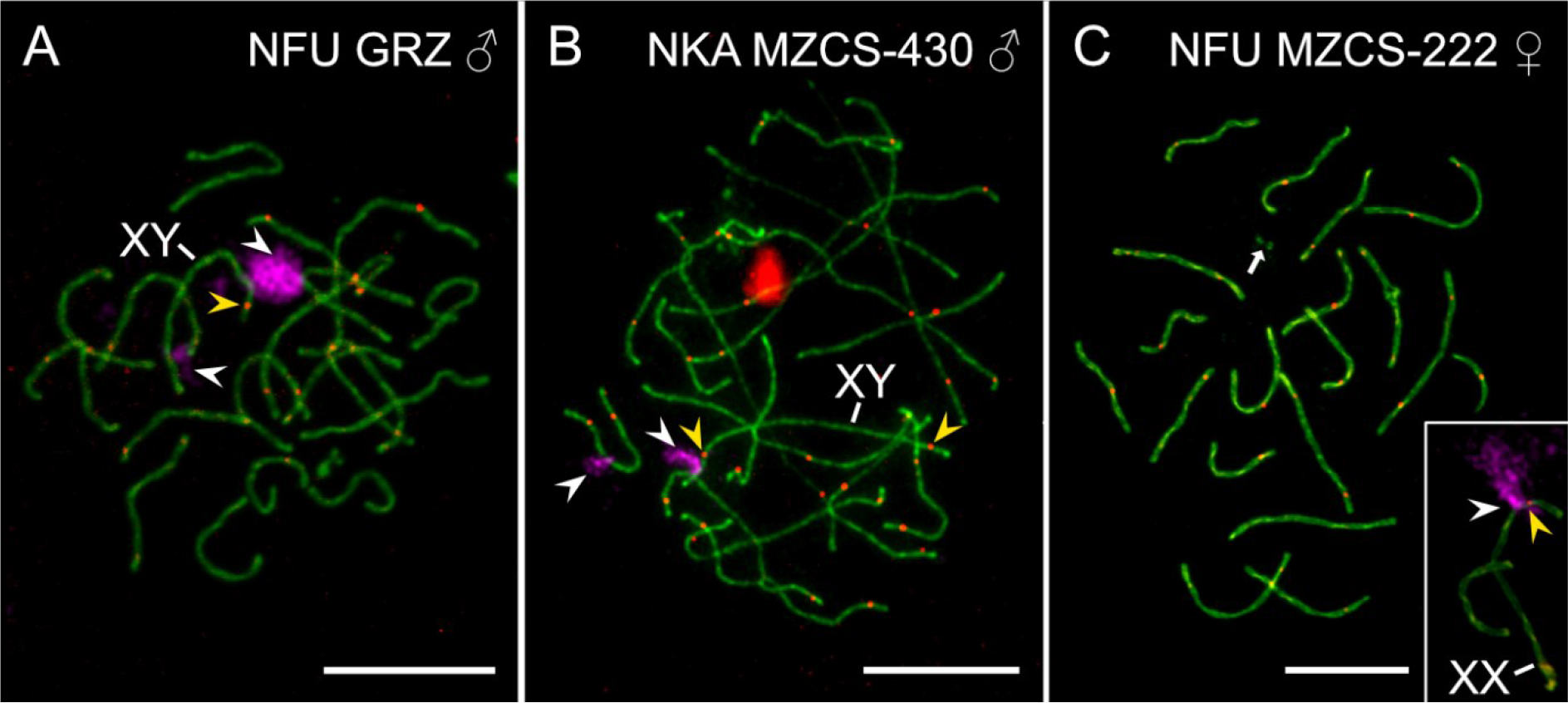
Selected *N. furzeri* and *N. kadleci* pachytene spreads after synaptonemal complex analysis and subsequent 18S rDNA FISH. (A,B) males; (C) female. SCs were visualized by anti-SYCP3 antibodies (green), the recombination sites were identified by anti-MLH1 antibodies (red). Slides were reprobed with 18S rDNA (magenta signals, white arrowheads). Based on patterns inferred from mitotic metaphases (see S8-S12 Figs), XY or XX bivalent is the larger from the two bivalents bearing 18S rDNA site. Yellow arrowheads point at MLH1 foci on XY or XX sex bivalent. Note that in analysed males both autosomes and sex chromosomes form standard bivalents without any irregularities in pairing. The sex chromosome bivalent contains one or two terminal/subterminal MLH1 foci. Full set of male plates from all studied populations is provided in S13 Fig. Female spreads studied in *N. furzeri* population MZCS-222 only (**C**) also displayed 19 standard bivalents, however, note an additional miniature bivalent-like object (arrow) which was present in virtually all analysed female spreads. Inset: XX bivalent from another spread, after reprobing with 18S rDNA probe. Note a single subterminal MLH1 focus adjacent to 18S rDNA site. Scale bar = 10 μm.

We were able to identify XY and XX sex bivalent from the complement based on sequential mapping of 18S rDNA probe onto the same meiocytes. Based on rDNA distribution patterns revealed on mitotic metaphases (Figs 2, S8, S9, S10, S11 and S12), we could infer that from two bivalents bearing the 18S rDNA signal the larger one was formed by XY sex chromosomes. The location of 18S rDNA signal further helped us to identify q- and p-arms of these chromosomes. In females, the number of analysable meiocytes after rDNA FISH was considerably reduced by prevailing high fluorescent background caused probably by high amount of transcripts and proteins in the cell. On the XY sex chromosome bivalent, often a single MLH1 signal was observed in a subtelomeric region on the q-arms only. Occasionally, a second signal was observed on the p-arms of the sex bivalent, adjacent to rDNA site. On the XX bivalent (studied only in females of *N. furzeri* population MZCS-222), the MLH1 foci distribution was highly similar, with a single MLH1 site adjacent to rDNA cluster on the p-arms. Finally, we tracked the possible signs of synaptic adjustment by FISH with telomeric probe (S14 Fig). The rationale is that sex chromosomes of different sizes should display different locations of telomeric signals if they are heteromorphic and aligned into bivalent without a synaptic adjustment (e.g. [59, 60]). In our sampling the telomeric signals were always confined to the very ends of all bivalents (S14 Fig), suggesting synaptic adjustment in heteromorphic XY bivalent.

## Discussion

Here, we show that *N. furzeri* and *N. kadleci* share an XY sex chromosome system. While in *N. furzeri*, our data corroborate previous findings in several populations including the GRZ strain [35,36,39], in *N. kadleci* we describe this sex chromosome system for the first time. Sex chromosomes in studied populations of both sister species are mostly heteromorphic (contrary to previous view based on genomics data [35]); and display highly similar patterns of differentiation as well as a putative MSD gene. Our BAC-FISH results strongly suggest that XY sex chromosomes of both species are homeologous which means that this sex chromosome system is evolutionarily older then previously anticipated. Sex chromosomes have not been previously identified in either of the studied species by means of conventional cytogenetics [44]. Here, we discuss only sex chromosome differentiation and evolution, while other findings (repeat landscape at karyotype level) are discussed in S3 Text.

First, we performed comparative genomic hybridization (CGH) to identify sex chromosome pair and assess its differentiation. CGH is based on competitive hybridization of male and female genomic probes and thus it can identify sequences unique to or enriched on a sex-limited chromosome. The method was used in different fish species with varied success (e.g. [19,55,56,61]). CGH highlighted centromeric regions but did not reveal any chromosomal segments labelled preferably by male genomic probe in *N. furzeri* and *N. kadleci* populations (Figs 2, S8 and S10). However, CGH detected a single chromosome with largely reduced hybridization signals in the centromeric region in all populations but NFU MZCS-121. Sequential analysis involving physical mapping of sex-linked BACs [35] confirmed it is the Y chromosome (Figs 2, S8 and S10). The deletion of a repetitive DNA segment from the Y contributed to the different size of X and Y chromosomes and might have resulted from intrachromosomal ectopic recombinations between centromeric repeats [62, 63].

To further examine differences between X and Y chromosomes, we performed the RepeatExplorer2 analysis of the repeatome in both *Notobranchius* spp. under study. We have focused on tandem repeats as these were misrepresented in the reference genome sequence of *N. furzeri* [35]. Three satellites, namely Nfu-SatA, Nfu-SatB, and Nfu-SatC, were identified among the most abundant repeat clusters comprising ≥ 0.5% genome in both *N. furzeri* and *N. kadleci* (Table 1). The Nfu-SatA and Nfu-SatB were previously characterized in more detail in (i) *N. furzeri* [35, 45]. FISH mapping of these satellites confirmed that they formed clusters overlapping with blocks of constitutive heterochromatin revealed by C-banding in all centromeres (cf. S3 and S5 Figs) and their distribution on sex chromosomes largely coincided with CGH hybridization signals. Nfu-SatA and Nfu-SatB were thus comprised in the region deleted from Y chromosomes (cf. S5 Fig with S8 and S10 Figs). The prevalence of Nfu-SatA and Nfu-SatB repeats in centromeric regions indicated that they play an important role in centromere organization and/or function in both killifish species examined. It has been hypothesized that asymmetry of female meiosis drives changes in copy number of centromeric repeats to ensure transfer of a chromosome into an oocyte [64, 65]. The observed reduction in centromeric cluster of the Nfu-SatA and Nfu-SatB repeats on Y chromosomes thus could reflect relaxed selection imposed by this so-called centromeric drive, as the Y chromosome never passes through females [66, 67].

The remaining tandem repeat, Nfu-SatC, differed substantially in its distribution patterns from the two centromeric satellites. Mapping of the Nfu-SatC revealed it was dispersed in small clusters across the entire genome in all studied strains of both *N. furzeri* and *N. kadleci* (S5 Fig). Yet, it formed a few pronounced accumulations with the most prominent on sex chromosomes, particularly on Y chromosomes, where it covered a large portion of long arms (Figs 2, S5, S8 and S10).

Indeed, the Nfu-SatC corresponds to the 634 nt tandem repeat, which forms a 35 Kb long cluster in the SD region in the MZM-0403 *N. furzeri* population [35] and it clearly spread on the q-arms of Y chromosomes of all studied strains but NFU MZCS-121. The size and location of Nfu-SatC are consistent across both studied species with the only exception of the NFU MZCS-121 population (Figs 2, 4 and S8). Accumulation of Nfu-SatC on Y chromosomes was probably facilitated by a low level of recombination along the q-arms of the Y chromosome as repetitive DNA in general populates these regions due to relaxed purifying selection [68–70]. Several repetitive DNA classes including microsatellites and satDNA have been found to accumulate in region of divergence on heterogametic sex chromosome in fishes [19,71–75]. Considering the extent of non-recombining region in the *N. furzeri* GRZ strain [35], we cannot exclude that the spread of the Nfu-SatC on Y chromosomes was further augmented by large inversions, which occurred independently in *N. furzeri* Chefu populations [35] as well as in *N. kadleci* (this study, Figs 2, 4, S7 and S10).

**Fig 4.**
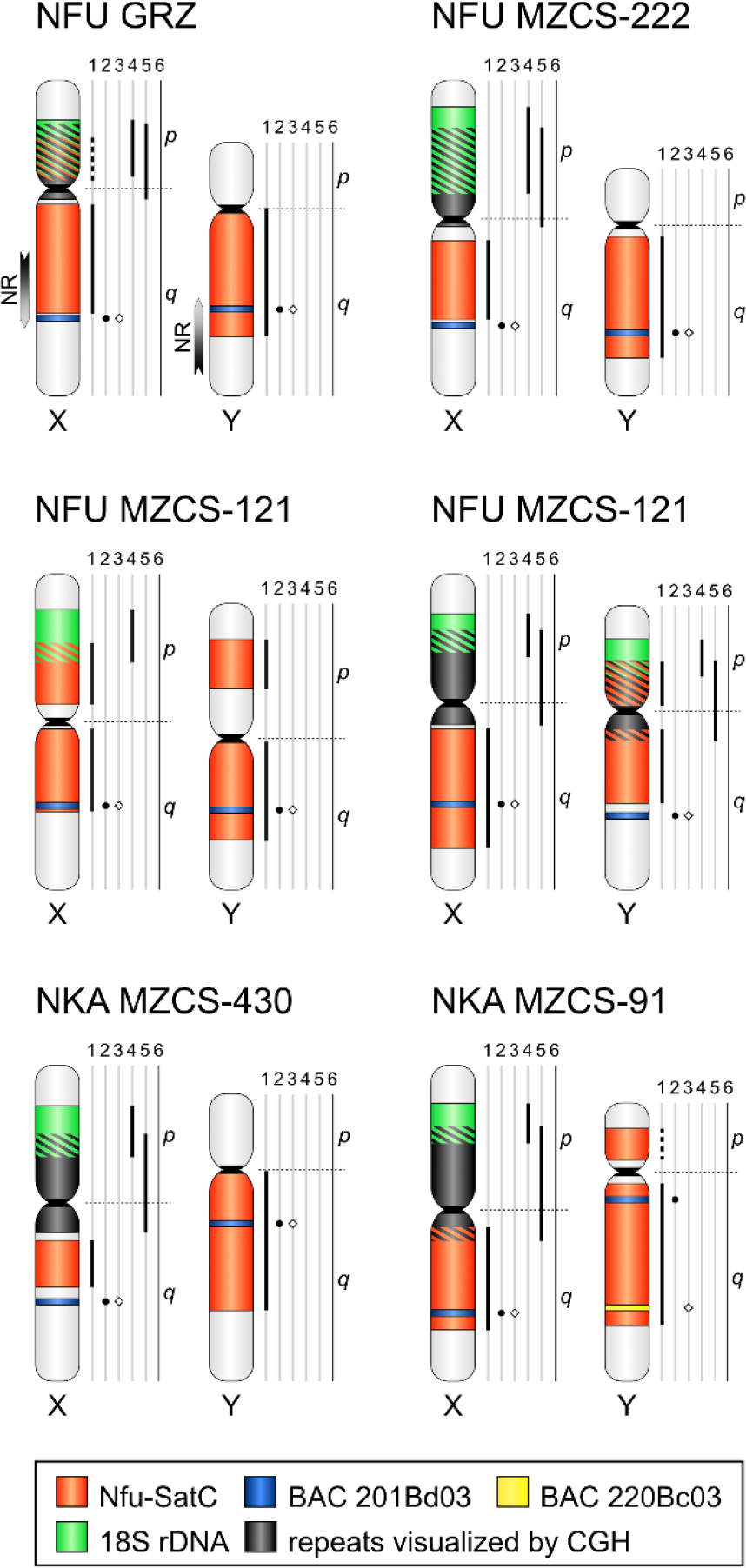
Integrative cytogenetic maps of XY sex chromosomes of studied *N. furzeri* and *N. kadleci* populations. Locations of BAC clones 201Bd03 (*gdf6*-bearing) and 220Bc03, 18S rDNA, Nfu-SatC satellite DNA and centromeric repeats revealed by CGH are depicted. All maps were constructed based metric analysis (details and data are provided in S4 Text including three associated tables, and in the Dryad digital data repository, doi: 10.5061/dryad.4mw6m90ck). Numbered vertical lines next to each chromosome depict the chromosomal location and size of the analysed markers in the following order: 1 – Nfu-SatC, 2 – BAC clone 201Bd03, 3 – BAC clone 220Bc03, 4 – 18S rDNA, 5 – CGH. Line 6 delineates short (p) arm and long (q) arm. “Empty” vertical lines mean: i) missing data (CGH repeats; one NFU3 MZCS-121 XY haplotype), ii) undetermined values (CGH repeats, all remaining Y chromosomes; not meassured due to their size reduction and restriction to the centromeric region), iii) absence of a given marker on the chromosome (remaining cases). Dashed horizontal line indicates the position of the centromere. Dashed vertical line in NFU GRZ and NKA MZCS-91 indicates the site with a presence/absence polymorphism. Due to their colocalization, the two mapped BAC clones were anchored by 201Bd03 and depicted as blue band in all populations but NKA MZCS-91. In the latter the clones were separated by an inversion. In NFU GRZ, the non-recombining region (NR) corresponding to 26 Mbp long inversion was delimited based on position of the 201Bd03 BAC bearing *gdf6* and the information from Reichwald et al. [35]. Note that *N. furzeri* males from Chefu clade populations (MZCS-222 and GRZ) feature very similar patterns of XY differentiation. In both populations XY sex chromosomes are heteromorphic, where Y chromosome is smaller than X counterpart and lacks block of repeats on its short arms. *N. furzeri* males from Limpopo clade (MZCS-121 population) display intra-population polymorphism in sex chromosome differentiation. Two Y haplotypes are shown. Note the altered sizes of Nfu-SatC on the q-arms of X and Y chromosomes when both XY variants in this population are compared. Males from both *N. kadleci* populations possess XY sex chromosome pair homeologous to the *N. furzeri* one, with pattern of differentiation similar to the Chefu clade. They also feature large Y-linked inversions, which differ in size between the two populations. Note that putative sex-determining allele of *gdf6* gene has been shifted towards centromere in both *N. kadleci* populations.

Differences between X and Y chromosomes were further confirmed by location of genes for major ribosomal RNA detected by FISH with the 18S rDNA probe. In most studied populations, the 18S rDNA locus was localized on the p-arms of the X chromosome while it was missing on the Y chromosome (Figs 2, S9, S10, S11 and S12). Nonetheless, intrapopulation polymorphisms were detected in NFU MZCS-121 and NKA MZCS-91 (Figs 2, S8-S2). In NFU MZCS-121, the Y chromosome of some males bore a rDNA locus, while none was detected on the X chromosome of some females. In contrast, polymorphic rDNA cluster on a single X chromosome and its total absence on the X chromosomes in one female was observed in NKA MZCS-91 (S10 and S12 Figs).

Altogether, our analyses of the distribution of various repetitive sequences clearly demonstrated that *N. furzeri* and *N. kadleci* sex chromosomes are heteromorphic. Moreover, as the pattern of their differentiation is similar between the Chefu populations of *N. furzeri* and its sister species *N. kadleci*, the changes are either convergent or occurred in a common ancestor of both species. Anyway, it seems the differentiation is not a result of inversions which rose independently in the Chefu populations of *N. furzeri* [35] and its sister species *N. kadleci* (Figs 2, 4, 5, S8 and S11).

Sex chromosome differentiation gradually affects their pairing in meiosis, regions with substantial sequence divergence might exhibit delayed pairing or remain asynaptic. Bivalent formed by heteromorphic sex chromosomes often undergoes size equalization also known as synaptic adjustment, as also reported from fishes [55,60,76,77]. Our analyses of meiotic chromosomes showed that despite their differentiation, X and Y sex chromosomes fully pair in *N. furzeri* and *N. kadleci*. Only occasionally, a putative unpaired or self-paired region was recorded in *N. furzeri* (S13B Fig). The sex chromosome also differed in size, yet standard telomeric FISH patterns indicated synaptic adjustment (S15 Fig). Immunostaining of MLH1 foci revealed either one (interstitial or terminal/subterminal) or two (terminal/subterminal) chiasmata in females. Bivalents with more chiasmata were rare (Fig 3C). In male meiosis, the patterns were similar (Figs 3 and S13). As for the sex chromosomes, there was usually only one chiasma between X and Y in the distal region of the q-arms (Fig 3; S14 Fig), although the second chiasma was also rarely detected on the p-arms of the XY bivalent, close to rDNA region. The sex chromosomes could have evolved either from a chromosome pair with reduced recombination or the recombination could have been reduced upon acquisition of the *gdf6Y* gene for sex determination (cf. [10,11,78]). Jeffries et al. [10] proposed that recombination between sex chromosomes can cease as a result of neutral accumulation of sequence divergence linked to the sex determining gene. Sex chromosome differentiation could have been further enhanced by heterochiasmy, i.e. sex-specific pattern of recombination. In fishes, recombination is mostly restricted to chromosomal ends in males and more evenly distributed across female chromosomes [79–82]. Heterochiasmy could have reduced recombination along the majority of the Y chromosomes once the *gdf6Y* gene took over sex determination (cf. [83]) as the Y chromosome is transmitted only through the paternal lineage. Thus, it would be sheltered from meiotic recombination, which could drive a convergent differentiation process even in the absence of inversions, which is in line with the alternative models [10,11,83]. We tried to compare the distribution of MLH1 foci between sexes, yet our analysis does not have sufficient power to test for heterochiasmy and further study of recombination landscape in the killifishes is much needed.

Reichwald et al. [35] hypothesized that the primary suppression of recombination occurred in the 196 Kb long region containing the putative MSD gene *gdf6Y* in *N. furzeri* ca. 750 thousand years ago (Kya) and that it was crucial for its fixation. However, as mentioned above, physical mapping of previously identified sex-linked BACs revealed that heteromorphic sex chromosomes of *N. furzeri* and *N. kadleci* are homeologous. The sex chromosome system thus evolved in a common ancestor of *N. furzeri* and *N. kadleci* upon their split from *N. orthonotus* and it is thus 2-5× older than previously assumed (approximately 1.5-4 My; cf. [35,41,46,84]). However, we cannot exclude that the killifish XY sex chromosome system evolved much earlier, as the system could be shared with *Fundulosoma thierryi* with X1X2Y multiple sex chromosome system formed via sex chromosome-autosome fusion [44] which diverged from *Nothobranchius* spp. ca. 14 Mya [41]. Whether the XY sex chromosomes of *N. furzeri* and *N. kadleci* share homology with *F. thierryi* remains to be tested.

Interestingly, while both *N. furzeri* populations from Chefu clade (MZCS-222 and GRZ) display highly similar patterns of XY differentiation, a single population from Limpopo clade (MZCS-121) features intra-population variability in terms of the Y chromosome differentiation (Fig 4). The primary non-recombining SD region was characterized in the *N. furzeri* MZM- 0403 population [35], which belongs to the Limpopo clade [31]. In our sampling, the clade is represented by the NFU MZCS-121 population, which differed from the others in the distribution of all studied repetitive sequences (see above and Figs 2 and 5). We hypothesize that the primary 196 Kb long SD region represents a linkage disequilibrium maintained by strong selection. In NFU MZCS-121 population, rather homomorphic XY might still occasionally recombine, and these rare events might translocate the primary SD region to the X chromosome, which resets the differentiation of the Y chromosome. The polymorphic distribution of Nfu-SatC and rDNA on both arms of the X and Y sex chromosomes (Figs 2, 4 and S8) fits this scenario. Alternatively, the recombination could have been restored by selection due to mutation load (cf. [12]).

**Fig 5.**
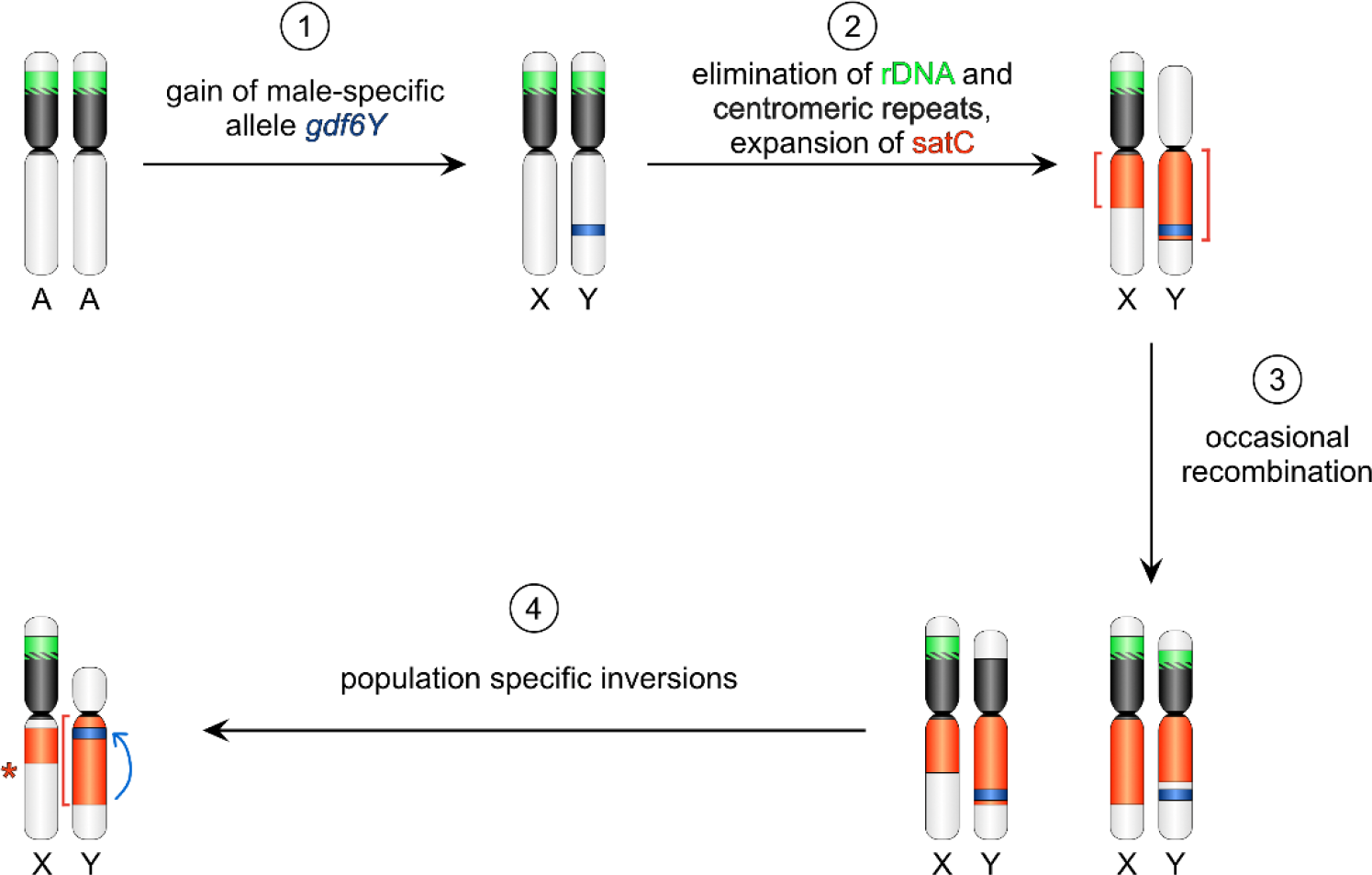
Scheme of differentiation of the Y chromosome in *N. furzeri* and *N. kadleci*. Upon *gdf6Y* aquired the role in sex determination (1), Y chromosome repetitive landscape diverged from its counterpart (2) presumably due to reduced recombination either in a common ancestor of both species or convergently in various populations. The recombination could have been suppressed following the neutral model by Jeffries et al. [10]. Yet the occasional recombination may still occur (3) as evidenced by mixed patterns observed in the *N. furzeri* Limpopo population MZCS-121. Eventually, secondary inversions occured (4) and the Y chromosome heteromorphy is maintained. The inversions could evolve under the Lenormand and Roze [12] model, in which inversions are fixed by sexually antagonistic regulatory effects. In such case the Limpopo polymorphism could represent restored recombination due to slow evolution of dosage compensation of degenerating genes within inversion.

Physical mapping of the *N. kadleci* sex chromosomes further revealed large polymorphic inversions analogous to those previously reported in *N. furzeri* populations [35]. This type of rearrangement obviously shifted the *gdf6* locus to the close proximity of centromere in males of both populations. Y-linked inversions thus seem to be common in annual *Nothobranchius* spp. and could be beneficial as they shelter the SD locus from occasional recombination (see above). The inversion might be achieved by intrachromosomal ectopic recombination between repeats [63, 85] and Nfu-SatC might provide a suitable substrate for this process. Reichwald et al. [35] hypothesized that the population-specific Y chromosome polymorphisms represent early stage of sex chromosome evolution preceding a selective sweep in which the most successful variant gets fixed across the species. Yet the polymorphic Y-linked inversions have been maintained for a long time. The age of the inversions was previously estimated at 38-70 Ky [35]. However, since the *N. furzeri* sex chromosome system is much older, the age of the inversions recalibrated according to Reichwald et al. [35] could reach up to 203-374 Ky. Absence of selective sweep thus suggests that formation of the Y-linked inversions in killifishes is not driven by SA selection despite their striking sexual dimorphism [86]. This agrees with autosomal linkage of male tail coloration identified in *N. furzeri* [87] and further supports our view that other alternative processes shape sex chromosome evolution in *N. furzeri* and *N. kadleci*. For instance, it has recently been proposed that Y-linked inversions could be maintained by early evolution of dosage compensation resulting in sexually antagonistic regulatory effects [12].

Studies of *Drosophila* neo-sex chromosomes showed that expression of protein coding genes is downregulated in 100 Ky old non-recombining region, while accumulation of non- sense mutations and repetitive sequences occurred ca. 1 My upon restriction of recombination [88]. In killifishes, it is reasonable to assume that loci outside the inversions still occasionally recombine as suggested by male single nucleotide variation analysed by Reichwald et al. [35]. Yet we observed dynamic changes in Y chromosome repetitive landscape which could be shaped by its recombination landscape or absence of centromeric drive and fixed by genetic drift (cf. [46]).

For further testing the above-mentioned hypotheses, *N. furzeri* populations from Limpopo clade seem to be especially informative due to lower degree of differentiation and higher level of polymorphisms. Besides that, the future aims will be i) to reveal the ancestral state of shared XY sex chromosome system, ii) whether the same sex chromosome system and putative MSD gene are driving sex determination also in other *Nothobranchius* species with so far undetermined sex chromosomes and iii) what is the evolutionary relationship of herein studied XY sex chromosome system to multiple sex chromosomes known in some *Nothobranchius* spp.

## Conclusions

To conclude, the *N. furzeri* and *N. kadleci* populations share basic karyotype patterns, the most abundant repeats and an XY sex chromosome system including the *gdf6Y* gene as putative MSD. The XY sex chromosomes are heteromorphic in the *N. furzeri* Chefu clade and *N. kadleci* and much older than previously assummed (approximately 1.7–4 Mya). In the Limpopo NFU MZCS-121 population, a polymorphism in the distribution of repeats suggests ongoing recombination between X and Y, which hinders the differentiation process. Despite their different size, meiotic analyses revealed almost no irregularities in X and Y chromosome pairing in the *N. furzeri* Chefu clade and *N. kadleci* populations implying synaptic adjustment of sex chromosomes. We propose that the observed differentiation is due to dynamic changes in repetitive DNA landscape outside inversions comprising MSD gene, which can be explained by non-canonical models of sex chromosome differentiation.

## Materials and Methods

### Fish sampling

We analysed individuals belonging to three populations of *N. furzeri* (two from Chefu clade and one from Limpopo clade) and two populations of *N. kadleci*. The detailed information is provided in Fig 1.

### Ethics statement

To prevent fish suffering, all handling of fish individuals followed European standards in agreement with §17 of the Act No. 246/1992 coll. The procedures involving fishes were supervised by the Institutional Animal Care and Use Committee of the Institute of Animal Physiology and Genetics CAS, v.v.i., the supervisor’s permit number CZ 02361 certified and issued by the Ministry of Agriculture of the Czech Republic. For direct preparations of chromosomes from kidney and gonads, fishes were euthanized using 2-phenoxyethanol (Sigma-Aldrich, St. Louis, MO, USA) before organ sampling. Fin samples (a narrow stripe of the tail fin) were taken from live individuals after fishes were anestheticized using MS-222 (Merck KGaA, Darmstadt, Germany).

### Conventional cytogenetics and chromosome banding

Mitotic or meiotic chromosome spreads were obtained either from regenerating caudal fin tissue as described by Völker and Ráb [89], with the modifications of Sember et al. [90] and altered time of fin regeneration (one week), or by direct preparation from the cephalic kidney and gonads [91]. In the latter, the quality of chromosomal spreading was enhanced by a dropping technique described by Bertollo et al. [92]. Chromosomes were either stained with 5% Giemsa solution (pH 6.8) (Merck, Darmstadt, Germany) for a conventional cytogenetic analysis, or left unstained for other methods. For fluorescence *in situ* hybridization (FISH), slides were dehydrated in an ascending ethanol series (70%, 80% and 96%, 3 min each) and stored in a freezer (-20 °C) until further use.

Constitutive heterochromatin distribution was analysed using C-banding [93]. Chromosomes were stained with 4′,6-diamidino-2-phenolindole (DAPI), 1.5 µg/mL in antifade, (Cambio, Cambridge, United Kingdom) and the pictures inverted for better contrast. Fluorescence staining was done by GC-specific fluorochrome Chromomycin A3 (CMA3) and AT-specific fluorochrome DAPI (both Sigma-Aldrich), following Mayr et al. [94] and Sola et al. [95].

### DNA isolation

High-weight molecular genomic DNA (gDNA) from male and female *N. furzeri* and *N. kadleci* individuals was extracted from liver, spleen and brain tissue using MagAttract HMW DNA Kit (Qiagen, Hilden, Germany) following the manufacturer’s instructions but with doubled volume of reagents and buffers except to elution buffer. gDNA was used for molecular cytogenetic applications or low-pass genome sequencing.

### Short-read genome sequencing

gDNA from one male and three females of *N. kadleci* (population MZCS-430) was randomly sheared and used to prepare Illumina paired-end libraries with 450 bp inserts at Novogene (HK) Co, Ltd. (Hong Kong, China). The libraries were sequenced on the Illumina NovaSeq 6000 platform using a protocol generating 150 bp paired-end reads with genome coverage at least 2.5×.

### Repetitive DNA analysis by RepeatExplorer2 pipeline

For the repeatome analysis, we used available datasets from GRZ strain of *N. furzeri* 35,36]; accession numbers ERR583468, ERR583470, ERR583471, SRR1261480) and de novo sequenced data from *N. kadleci*. Repetitive sequences were characterized directly from quality-filtered reads using a graph-based sequence clustering method implemented in the RepeatExplorer2 software package. The pipeline also allows the estimation of repeat quantities from number of reads in clusters and their annotation [52].

First, the quality of raw data was checked using FastQC (version 0.11.5; [96]) and all datasets were filtered with Cutadapt (version 1.15; [97] using the following parameters: “-- nextseq-trim=20 --length 100 --m 100 -M 100 -a AATGATACGGCGACCACCGAGATCTACACTCTTTCCCTACACGACGCTCTTCCGA TCT -A GATCGGAAGAGCACACGTCTGAACTCCAGTCACNNNNNNATCTCGTATGCCGTCT TCTGCTTG”. Resulting reads of 100 bp length were converted to fasta format, interlaced and subsampled with the RepeatExplorer2 suite in the command line. For detailed study of the repetitive landscape within species, 1,187,000 reads were pseudorandomly subsampled from three female datasets, in total representing ca. 0.375× coverage of the genome. Finalized datasets were processed via Galaxy RepeatExplorer portal (https://repeatexplorer-elixir.cerit-sc.cz/galaxy/). RepeatExplorer2 clustering was performed with settings for comparative analysis, using Metazoa v. 3.0 reference database and automatic filtering of abundant satellite repeats.

### Repeat annotation

Annotation of individual clusters is based on RepeatExplorer2 and TAREAN automatic annotations as well as RepeatMasker (https://www.repeatmasker.org/) [98] protein-based masking of contigs corresponding to respective clusters. Putative satellites were annotated based on small dense globular shape of graph, which is usually typical for this type of repeat and/or tandemly repeated sequence in at least one contig, produced by RepeatExplorer2.

To identify repeats homologous between species, *N. kadleci* contigs corresponding to RepeatExplorer2 clusters of interest were searched against *N. furzeri* RepeatExplorer2 database using BLAST/N. Only matches with e-value lower than 10^-5^, longer than 200 bp, and similarity higher than 85% were considered.

### Primer design

To amplify selected repetitive DNA elements, custom primers were designed in Geneious Prime 2020.1.2 (https://www.geneious.com). Primer sequences are listed in S1 Table.

### Preparation of FISH probes

Amplification of 5S and 18S rDNA and two satellite DNA motifs from RepeatExplorer analysis was done by polymerase chain reaction (PCR) with primers and thermal profiles specified in S1 and S2 Tables, respectively. The obtained PCR products were purified using NucleoSpin Gel and PCR Clean-up (Macherey-Nagel GmbH, Düren, Germany) according to supplier’s instructions. Amplification products with multiple bands were electrophoresed in 0.8% agarose gels and each fragment was purified using the same kit as described above. The subsequent procedures involving cloning of the purified products, plasmid isolation, sequencing (in both strands) of selected positive clones, assembly of chromatograms from obtained sequences and sequence alignment followed essentially the same workflow as described in Sember et al. [99]. The custom sequencing was done in Macrogen company (Netherlands). The content of resulting consensus sequences was verified using NCBI BLAST/N analysis [100] and selected clones were used for a FISH probe preparation. In case of chromosomal mapping of 18S rDNA, for most experiments we used the already optimized probe obtained from a bighead carp *Hypophthalmichthys molitrix* (for details, see 54]).

rDNA probes were labelled either by nick translation (18S rDNA) or by PCR (5S rDNA) following the conditions and set of primers provided in S1 and S2 Tables. The labelling nucleotide mix contained either biotin-16-dUTP or digoxigenin-11-dUTP (both Roche, Mannheim, Germany). Due to its long size, the 18S rDNA probe was generated in two steps:

(i) non-labelling PCR amplification from a verified 18S rDNA clone and (ii) nick translation (1 h 40 min) of the amplified 18S rDNA product using Nick Translation Mix (Abbott Molecular, Illinois, USA). In case of two satellite DNAs retrieved from RepeatExplorer, namely Nfu-SatB and Nfu-SatC (see Results section), probes were labelled by nick translation with Cy3-dUTP using Cy3 NT Labeling Kit (Jena Bioscience, Jena, Germany). In this case, the entire plasmid DNA was used as template. The optimal fragment size of the probe (ca. 200–500 bp) was achieved after 30 min of incubation at 15 °C.

A dual-color rDNA FISH for each slide involved 250 ng of each probe and 25 µg of sonicated salmon sperm DNA (Sigma-Aldrich). In case of single-colour FISH with satDNA, 250-500 ng of nick-translated plasmid was used per slide, supplemented with 12.5-25 µg of sonicated salmon sperm DNA (Sigma-Aldrich). For 18S rDNA FISH on synaptonemal complexes, 500 ng of the probe and 25 µg of sonicated salmon sperm DNA (Sigma-Aldrich) per slide was used. The final hybridization mixtures for each slide (15 µL) were prepared according to Sember et al. [90].

### BAC clones’ isolation and probe preparation

For sex chromosome identification, we chose four BAC clones carrying X and/or Y linked sequences from genomic library of *N. furzeri* ([35]; S3 Table) and mapped them chromosomally by BAC-FISH. The selected BAC clones were identified in the BAC library thanks to their identification codes, isolated using NucleoBond Xtra Midi kit (Macherey Nagel) and verified by PCR using primers specified in S4 Table. BAC DNA was labelled following the protocol of Kato et al. [101] with modifications described in Yoshido et al. [102]. For each dual-colour FISH experiment, the final probe mixture contained 300 ng of Cy3-labelled BAC and 500 ng of FITC-labelled BAC. For blocking unspecific hybridization, 6 µg of thermally fragmented male gDNA (99 °C, 20 min), 1500 ng of degenerate oligonucleotide-primed (DOP) PCR-derived competitive DNA from male gDNA (based on Kosyakova et al. [103]) and 25 µg of sonicated salmon sperm DNA (Sigma-Aldrich) were added. The probe was ethanol-precipitated and dissolved in 10 µL of hybridization mixture containing 50% deionized formamide (Carl Roth GmbH, Karlsruhe, Germany) and 10% dextran sulfate (Sigma-Aldrich).

### Probes for comparative genomic hybridization (CGH)

We employed CGH for identification of Y sex chromosomes as this method may help to delimit roughly the region of differentiation on heterogametic sex chromosome if this process already proceeded to certain stage of repetitive DNA accumulation. Simultaneous mapping of whole genome-derived probes from male and female (differentially labelled) to male chromosome spreads may thus reveal regions with biased or exclusive hybridization of male genomic probe (e.g. [55, 56]). For each population of *N. furzeri* and *N. kadleci*, male and female gDNA was labelled by nick translation with Cy3 NT Labeling Kit and Fluorescein NT Labeling Kit (Jena Bioscience), respectively. The optimal range of labelled DNA fragment sizes was achieved after 30 min of incubation at 15 °C. To block the excess of shared repetitive sequences we used unlabelled competitive DNA prepared from female gDNA by DOP-PCR (see above). The ratio of the probe vs. blocking DNA (see below) was chosen based on previous experience with the CGH method in other fishes (e.g. [19,56,61]). The final probe mixture for each slide was prepared by mixing male and female genomic probes (500 ng each), 6 μg of female-derived competitive DNA and 50 µg of sonicated salmon sperm DNA (Sigma-Aldrich). Each ethanol-precipitated probe mixture was dissolved in 20 μL of the hybridization buffer (for composition, see [90]).

### FISH analysis

The FISH experiments were done using a combination of two previously published protocols [61, 90], with slight modifications. Briefly, the preparations were thermally aged overnight at 37 °C and then 60 min at 60 °C, followed by treatments with RNase A (200 µg/mL in 2× SSC, 60–90 min, 37 °C) (Sigma-Aldrich) and then pepsin (50 µg/mL in 10 mM HCl, 3 min, 37 °C). In case of CGH and BAC-FISH experiments, the slides were subsequently incubated in 1% formaldehyde in 1× PBS (10 min) to stabilize the chromatin structure. For recycled slides subjected to reprobing in sequential experiments, the 1% formaldehyde treatment directly followed after the washing process described below (see section “Slide treatment for reprobing”). Denaturation of chromosomes was done in 75% formamide in 2× SSC (pH 7.0) (Sigma-Aldrich) at 72 °C, for 3 min. The hybridization mixture was denatured for 6 min (repetitive DNA probes) or 8 min (CGH) at 86 °C, or for 5 min at 90 °C (BAC-FISH). In case of CGH, the probes were pre-hybridized at 37 °C for 45 min to outcompete the repetitive fraction. After application of the probe cocktail on the slide, the hybridization took place in a moist chamber at 37 °C for 24 h (repetitive DNA probes) or 72 h (CGH and BAC-FISH). Subsequently, non-specific hybridization was removed by post-hybridization washes which followed different protocols based on the type of probe labelling.

In the case of fluorochrome-labelled probes, following Yano et al. [61] the preparations were treated two times in 1× SSC (pH 7.0) (65 °C, 5 min each) and once in 4× SSC in 0.01% Tween 20 (42 °C, 5 min). Slides were then washed in 1× PBS (1 min), passed through an ethanol row and mounted in antifade containing 1.5 µg/mL DAPI (Cambio, Cambridge, United Kingdom). Only in the case of BAC-FISH experiments, the post-hybridization washes followed different protocol [102]. Specifically, slides were treated once in 0.1× SSC in 1% Triton X-100 (5 min at 62 °C) and then in 2× SSC in 1% Triton X-100 (2 min at RT). Final wash in 1% Kodak PhotoFlo in H2O (2 min at RT) was followed by ethanol row and counterstaining with 0.2 g/mL DAPI mounted in antifade based on DABCO (1,4-diazabicyclo(2.2.2)-octane; Sigma-Aldrich, St. Louis, MO, USA).

In the case of hapten-labelled probes, following Sember et al. [90] the preparations were treated two times in 50% formamide/2× SSC (42 °C, 10 min) and three times in 1× SSC (42 °C, 7 min). Prior to the probe detection, 3% bovine serum albumin (BSA) (Vector Labs, Burlington, Canada) in 0.01% Tween 20/4× SSC was applied to the slides to block unspecific binding of antibodies. Hybridization signals were detected by Anti-Digoxigenin-FITC (Roche; dilution 1:10 in 0.5% BSA/PBS) and Streptavidin-Cy3 (Invitrogen Life Technologies, San Diego, CA, USA; dilution 1:100 in 10% NGS (normal goat serum)/PBS). In case of rDNA dual-colour FISH, although experiments with both possible combinations of labelling of 18S and 5S rDNA were included (to verify the observed patterns), all FISH images presented here have a unified coloring system – red for the 5S rDNA and green for 18S rDNA probes. Finally, slides were counterstained with DAPI as described above.

### Telomeric FISH

Telomeric (TTAGGG)*n* repeats were detected by FISH using a commercial telomere PNA probe directly labelled with Cy3 (DAKO, Glostrup, Denmark) according to the manufacturer’s instructions, with a single modification concerning the prolonged hybridization time (1.5 h).

### Non-denaturing FISH (ND-FISH)

We followed the protocol of Cuadrado and Jouve [104] (formerly used for chromosomal mapping of simple sequence repeats), with several modifications.

A probe mixture for one slide (30 µL) contained 2 pmol/µL of Nfu-SatA (5’ labelled by Cy3) oligonucleotides in 2× SSC. The probe mixture was denatured at 80 ℃ for 5 min, placed on ice (15-20 min) and then transfered to chromosome preparations which did not undergo any pre-treatment step and even no denaturation. After hybridzation (2-3 h, RT) unspecific probe hybridization was removed by treatment in 4× SSC/0.2% Tween 20 (10 min) and 4× SSC/0.1% Tween (5 min). Chromosomes were counterstained with 20 µL DAPI as described above.

### Synaptonemal complex analysis by immunostaining

We performed immunodetection of synaptonemal complex protein SYCP3 and mismatch repair protein MLH1, a marker for recombination sites, on male and female pachytene spreads in order to analyse the chromosome pairing properties with emphasis on XY and XX sex bivalents. Synaptonemal complex spreads were prepared from testes or ovaries following the protocol for *Danio rerio* [105–107] with some modifications. Briefly, dissected testes were suspended in 200-600 µL (based on the testes size and the cell density) of cold PBS. One microlitre of cell suspension was added to each 30 µL drop of cold hypotonic solution (PBS:H2O, 1:2) and placed on poly-l-lysine dry slides. After 20 min (*N. furzeri*) or 25-30 min (*N. kadleci*), slides were fixed with freshly prepared cold 2% formaldehyde (pH 8.0 – 8.5, with 0.03% SDS) for 3 min. Slides were then washed three times in 0.1% Tween 20 (pH 8.0-8.5), 1 min each and were dried for 1 h. Afterwards, immunofluorescence analysis of synaptonemal complexes was performed, using antibodies against the proteins SYCP3 and MLH1. The primary antibodies – rabbit anti-SYCP3 (1:300; Abcam, Cambridge, UK; RRID: AB_301639) and mouse anti-MLH1 (1:50, Abcam; RRID: AB_300987) – were diluted (v/v) in 3% BSA in 0.05% Triton X-100/PBS and applied onto the slides. The incubation was carried out overnight in a humid chamber at 4 °C. Next day, slides were washed three times in 0.1% Tween 20 in PBS, 10 min each and secondary antibodies, diluted (v/v) in 3% BSA in 0.05% Triton X- 100/PBS, were applied. Specifically, we used goat anti-rabbit Alexa 488 (1:300; Abcam; RRID: AB_2630356) and goat anti-mouse Alexa555 (1:100; Abcam; RRID: AB_2687594) and the slides were incubated for 4 h at RT. Then, after washing in 0.1% Tween 20 in PBS (three times, 10 min each) and brief washing in 0.01% Tween 20 in distilled H2O, slides were mounted in antifade containing DAPI, as described above.

### Slide treatment for reprobing

Preparations subjected to repeated sequential experiments were soaked in distilled water for 10 min. Then, after nail polish removal, slides were incubated in 4× SSC/0.1% Tween 20 (37 °C for 30 min). After subsequent cover slip removal and another incubation in 4× SSC/0.1% Tween 20 (37 °C for 30 min) slides were passed through an ascending ethanol row and fixed in methanol:acetic acid solution (3:1; v/v) for 30 min. After thermal aging (37 °C for 2h), slides were postfixed in 1% formaldehyde and further treated according to subsequent FISH protocol.

Slides reused after synaptonemal complex analysis were soaked in distilled water for 10 min and then washed three times in 4× SSC/0.1% Tween 20 (RT, 10 min each; including cover slip removal). Slides were then passed through an ethanol series and after air drying, they were subjected to a chromosome denaturation step in subsequent FISH protocol.

Slides reused for repeated BAC-FISH experiments (i.e. for BAC clone physical map generation) were reprobed following Yoshido et al. [102].

### Microscopic analyses and image processing

Results from molecular cytogenetic protocols were inspected using BX53 Olympus microscope equipped with appropriate fluorescence filter set. Images were captured under immersion objective 100× with a black and white CCD camera (DP30W Olympus) for each fluorescent dye separately using DP Manager imaging software (Olympus). The same software was used to superimpose the digital images with the pseudocolours. The digital images were then pseudocoloured (blue for DAPI, red for Cy3, green for FITC) and composed images were then optimized and arranged using Adobe Photoshop, version CS6.

Giemsa-stained preparations were analysed under Axio Imager Z2 microscope (Zeiss, Oberkochen, Germany), equipped with automated metaphase search (Metafer-MSearch scanning platform). Metaphase images were captured under 100× objective using CoolCube 1 b/w digital camera (MetaSystems, Altlussheim, Germany). The karyotypes were arranged using Ikaros software (MetaSystems, Altlussheim, Germany).

At least 20 chromosome spreads per individual and method were analysed, some of them sequentially. Chromosomes were classified according to Levan et al. [108], but modified as m – metacentric, sm – submetacentric, st – subtelocentric, and a – acrocentric, where st and a chromosomes were scored together into st-a category. Chromosome pairs were arranged according to their size in each chromosome category. Final images were optimised and arranged using Adobe Photoshop, version CS6.

## Supporting information

**S1 Fig. Karyotypes of *N. furzeri* males and females after Giemsa staining.** m = metacentric chromosome, sm = submetacentric chromosome, st/a = subtelocentric-to- acrocentric chromosome.

(PDF)

**S2 Fig. Karyotypes of *N. kadleci* males and females after Giemsa staining.** m = metacentric chromosome, sm = submetacentric chromosome, st/a = subtelocentric-to- acrocentric chromosome.

(PDF)

**S3 Fig. Mitotic metaphases of *N. furzeri* and *N. kadleci* after C-banding.** Chromosomes stained with DAPI (inverted picture). Putative Y sex chromosomes are marked. Scale bar = 10 μm.

(PDF)

**S4 Fig. Mitotic metaphases of *N. furzeri* and *N. kadleci* after CMA3/DAPI staining.** For better contrast, images were pseudocoloured in red (for CMA3) and green (for DAPI). (**A**) X chromosome is marked based on morphology and strong CMA3-positive signal related to major rDNA site. Scale bar = 10 μm.

(PDF)

**S5 Fig. FISH with tandem repeats retrieved from RepeatExplorer analysis on male metaphases.** Red signals: (**A**-**E**) Nfu-SatA minisatellite, (**F**-**J**) Nfu-SatB satDNA, (**K**-**O**) Nfu- SatC satDNA. Yellow signals: FISH with two differently labeled Nfu-SatC probes derived from cloned fragments with different sequence compositions (sequences were deposited in GenBank under accession numbers OM542182 and OM542183); note that green and red signals are entirely overlapping, producing virtually uniform yellowish hybridization pattern. Y sex chromosomes are marked if possible. Asterisks point to population-specific gap in signal in one metacentric pair in NFU MZCS-121 (**H**, **M**). Chromosomes were counterstained with DAPI (blue). Scale bar = 10 μm.

(PDF)

**S6 Fig. PNA-FISH with telomeric probe**. For better contrast, pictures were pseudocoloured in green (telomeric repeat probe) and red (DAPI). Scale bar = 10 μm.

(PDF)

**S7 Fig.** Physical map of selected XY-linked BAC clones. Details on BAC clones are provided in S3 and S4 Tables. BAC bearing *gdf6* gene is 201Bd03. Note that in NKA MZCS-430 male (**D**), three out of four BACs less intense and diffuse signals on the Y chromosome, probably due to sequence divergence on this sex chromosome. In females of this population (I), we did not get reproducible signals of the 262Ad08 clone, however, its position is known from male X chromosome. Note the shift in position of all four BACs (201Bd03, 220Bc03, 225De03 and 262Ad08) on the Y chromosome of NKA MZCS-430 male, while only three BACs (201Bd03, 225De03 and 262Ad08) are shifted in NKA MZCS-91 male. (**K**) Detailed scheme showing this inversion polymorphism in *N. kadleci* narrowed to two informative BACs.

(PDF)

S8 Fig. Sequential FISH experiments in N. furzeri males – full karyotypes.

The same metaphase plates were treated subsequently with different FISH protocols. i) BAC-FISH – dual-colour FISH with X-linked 220Bc03 BAC clone (green signal) and Y-linked 201Bd03 BAC (gdf6y-bearing clone; red signal); ii) single-colour FISH with Nfu-SatC probe (red signals); iii) dual-colour FISH with 5S (red signals) and 18S (green signals) rDNA probes. Note that 18S rDNA cluster is present on short arms of X chromosome but mostly not on Y chromosome. Some males from Limpopo population NFU MZCS-121 have exceptionally 18S rDNA site also on Y chromosome. iv) comparative genomic hybridization (CGH) performed sequentially after BAC-FISH on different set of metaphases – hybridization patterns of male (red) and female (green) genomic probes yielded uniform yellowish signal. No exclusive or biased accumulation of either probe was observed, failing thus to unmask the region of differentiation on Y chromosome. Note, however, differences between X and Y in the size of (peri)centromeric block of accumulated repetitive DNA. m = metacentric chromosome, sm = submetacentric chromosome, st = subtelocentric chromosome, st/a = subtelocentric-to- acrocentric chromosome.

S9 Fig. Sequential FISH experiments in N. furzeri females – full karyotypes.

The same metaphase plates were treated subsequently with different FISH protocols. i) BAC-FISH – dual-colour FISH with X-linked 220Bc03 BAC clone (green signal) and Y-linked 201Bd03 BAC (gdf6y-bearing clone; red signal); ii) single-colour FISH with Nfu-SatC probe (red signals); iii) dual-colour FISH with 5S (red signals) and 18S (green signals) rDNA probes. Note that 18S rDNA cluster is present on short arms of X chromosomes but some females from Limpopo population NFU MZCS-121 lacked 18S rDNA site on one X homologue. m = metacentric chromosome, sm = submetacentric chromosome, st/a = subtelocentric-to- acrocentric chromosome.

S10 Fig. Sequential FISH experiments in N. kadleci males and females – full karyotypes.

The same metaphase plates were treated subsequently with different FISH protocols. i) BAC-FISH – dual-colour FISH with X-linked 220Bc03 BAC clone (green signal) and Y-linked 201Bd03 BAC (gdf6y-bearing clone; red signal); ii) single-colour FISH with Nfu-SatC probe (red signals); iii) dual-colour FISH with 5S (red signals) and 18S (green signals) rDNA probes. Note that 18S rDNA cluster is present on short arms of X chromosome but not on Y chromosome. iv) comparative genomic hybridization (CGH) performed sequentially after BAC-FISH on different set of metaphases – hybridization patterns of male (red) and female (green) genomic probes yielded uniform yellowish signal. No exclusive or biased accumulation of either probe was observed, failing thus to unmask the region of differentiation on Y chromosome. Note, however, differences between X and Y in the size of (peri)centromeric block of accumulated repetitive DNA. m = metacentric chromosome, sm = submetacentric chromosome, st = subtelocentric chromosome, st/a = subtelocentric-to-acrocentric chromosome.

(PDF)

**S11 Fig. Karyotypes of *N. furzeri* after 5S/18S rDNA FISH.** 5S rDNA (red, arrows) and 18S rDNA (green, arrowheads); probes mapped on mitotic chromosomes. Chromosomes were counterstained with DAPI (blue). Note the presence of 18S rDNA site on X chromosomes and in NFU MZCS-121 males also on Y chromosome. Inter-individual variability in 18S rDNA sites is boxed. m = metacentric chromosome, sm = submetacentric chromosome, st/a = subtelocentric-to-acrocentric chromosome. Scale bar = 10 μm.

(PDF)

**S12 Fig. Karyotypes of *N. kadleci* after 5S/18S rDNA FISH.** 5S rDNA (red, arrows) and 18S rDNA (green, arrowheads); probes mapped on mitotic chromosomes. Chromosomes were counterstained with DAPI (blue). Note the presence of 18S rDNA site on X chromosomes. Inter-individual variability in 18S rDNA sites is boxed. m = metacentric chromosome, sm = submetacentric chromosome, st/a = subtelocentric-to-acrocentric chromosome. Scale bar = 10 μm.

(PDF)

**S13 Fig. Male pachytene spreads after synaptonemal complex analysis and subsequent 18S rDNA FISH.** SCs were visualized by anti-SYCP3 antibodies (green), the recombination sites were identified by anti-MLH1 antibodies (red). Except for (**E**), slides were recycled for 18S rDNA FISH (magenta signals, white arrowheads). Based on patterns inferred from mitotic metaphases (see S8, S10-S12 Figs), XY bivalent is the larger from the two bivalents bearing 18S rDNA site. Yellow arrowheads point on MLH1 foci on XY sex bivalent. Empty yellow arrowheads denote putative MLH1 sites. Arrow (**B**) points on putative self-paired region found on XY bivalent based on 18S rDNA FISH confirmation (inset). Asterisk (**C**) points on highly probably false positive 18S rDNA signal (based on evidence on many meiotic and mitotic plates from individuals from NFU MZCS-121 population). Note that analysed males form 19 standard bivalents without asynapses, including XY sex bivalent, with a rare exception depicted in (**B**).

Scale bar = 10 μm.

(PDF)

**S14 Fig. Male pachytene spreads after synaptonemal complex analysis and subsequent telomeric and 18S rDNA FISH.** SCs were visualized by anti-SYCP3 antibodies (green). SYCP3 immunostaining was followed by telomeric PNA FISH (red signals) (**A-E**) and in some cases additionally by 18S rDNA FISH (magenta signals, arrowheads) (**B**,**D**,**E**). Based on patterns inferred from mitotic metaphases (see S8, S10-S12 Figs), XY bivalent is the larger from the two bivalents bearing 18S rDNA site. Note that all telomeric signals are placed at the very ends of all bivalents which in the case of XY sex bivalent suggests synaptic adjustment in populations with heteromorphic sex chromosomes (i.e. all but NFU MZCS-121 where XY is often homomorphic). Scale bar = 10 μm.

(PDF)

S1 Text. Characterization of major satellite DNA markers

(PDF)

S2 Text. Patterns of distribution of 18S rDNA

(PDF)

S3 Text. General discussion about observed repetitive DNA patterns in *N. furzeri* and *N. kadleci* karyotypes.

(PDF)

**S4 Text. Reconstruction of X and Y cytogenetic maps in *N. furzeri* and *N. kadleci* populations.** S4 Text – Table 1. Means of the relative length and position of four cytogenetic markers mapped on X and Y chromosomes from mitotic metaphases in *N. furzeri*. S4 Text – Table 2. Means of the relative length and position of five cytogenetic markers mapped on X and Y chromosomes from mitotic metaphases in *N. kadleci*. S4 File – Table 3. Means of the absolute lengths of sex chromosomes in µm. Mean values represent the % of the total length of the sex chromosome. Distance values: from the tip of the p-arm to the location of the marker; length values: from the tip to the end of the marker region. N – number of sex chromosomes measured; SD – standard deviation.

(PDF)

S1 Table. Primers used for repetitive DNA amplification in this study.

(PDF)

S2 Table. PCR thermal profiles used in this study.

(PDF)

S3 Table. Selected BAC clones (following information from Reichwald et al. [35], supplement S1I)

(PDF)

S4 Table. Primers for BAC clone verification

(PDF)

## Supporting information

Supporting material for Stundlova et al.

## Acknowledgments

The authors are grateful to C. Englert, S. Förste and M. Platzer (The Leibniz Institute on Aging – Fritz Lipmann Institute, Jena, Germany) for providing BAC clones from the *N. furzeri* genomic library. We are also thankful to P. Šejnohová for laboratory assistance.

## Author contributions

**Conceptualization:** Alexandr Sember, Petr Nguyen, Martin Reichard

**Data curation:** Petr Nguyen, Anna Voleníková, Martina Dalíková, Tomáš Dvořák **Formal analysis:** Jana Štundlová, Petr Nguyen, Anna Voleníková, Martina Dalíková **Funding acquisition:** Alexandr Sember

**Investigation:** Jana Štundlová, Monika Kreklová, Karolína Lukšíková, Anna Voleníková, Tomáš Pavlica, Marie Altmanová, Martina Dalíková, Šárka Pelikánová, Anatolie Marta, Sergey A. Simanovsky, Matyáš Hiřman, Marek Jankásek, Tomáš Dvořák, Joerg Bohlen, Petr Nguyen, Alexandr Sember

**Methodology:** Jana Štundlová, Alexandr Sember, Petr Nguyen, Anna Voleníková, Martina

Dalíková, Anatolie Marta, Sergey A. Simanovsky

**Project administration:** Alexandr Sember, Petr Nguyen

**Resources:** Alexandr Sember, Petr Nguyen, Martin Reichard, Petr Ráb

**Supervision:** Alexandr Sember, Petr Nguyen, Martin Reichard, Petr Ráb, Jörg Bohlen

**Validation:** Jana Štundlová, Alexandr Sember, Petr Nguyen

**Visualization:** Jana Štundlová, Marie Altmanová, Karolína Lukšíková, Jörg Bohlen

**Writing ± original draft:** Alexandr Sember

**Writing ± review & editing:** Petr Nguyen, Alexandr Sember, Jana Štundlová, Martin Reichard, Anna Voleníková, Marie Altmanová, Jörg Bohlen, Petr Ráb, Sergey A. Simanovsky, Martina Dalíková

## Data availability statement

The nucleotide sequences of rDNA genes were deposited in NCBI GenBank under the following accession numbers: MZ694985 for 5S rDNA from *N. kadleci* and MZ682111 for 18S rDNA from *N. guentheri*. Meassurements related to the construction of cytogenetic maps of sex chromosomes are available from the Dryad digital data repository: https://doi.org/10.5061/dryad.4mw6m90ck. All other relevant data are within the paper and its Supporting Information file.

