## Supporting material for Stundlova et al. for "Sex chromosome differentiation via changes in the Y chromosome repeat landscape in African annual killifishes *Nothobranchius furzeri* and *N. kadleci*"

**The PDF file includes:**

S1 Fig
S2 Fig
S3 Fig
S4 Fig
S5 Fig
S6 Fig
S7 Fig
S8 Fig
S9 Fig
S10 Fig
S11 Fig
S12 Fig
S13 Fig
S14 Fig
S1 Text
S2 Text
S3 Text
S4 Text
S1 Table
S2 Table
S3 Table
S4 Table

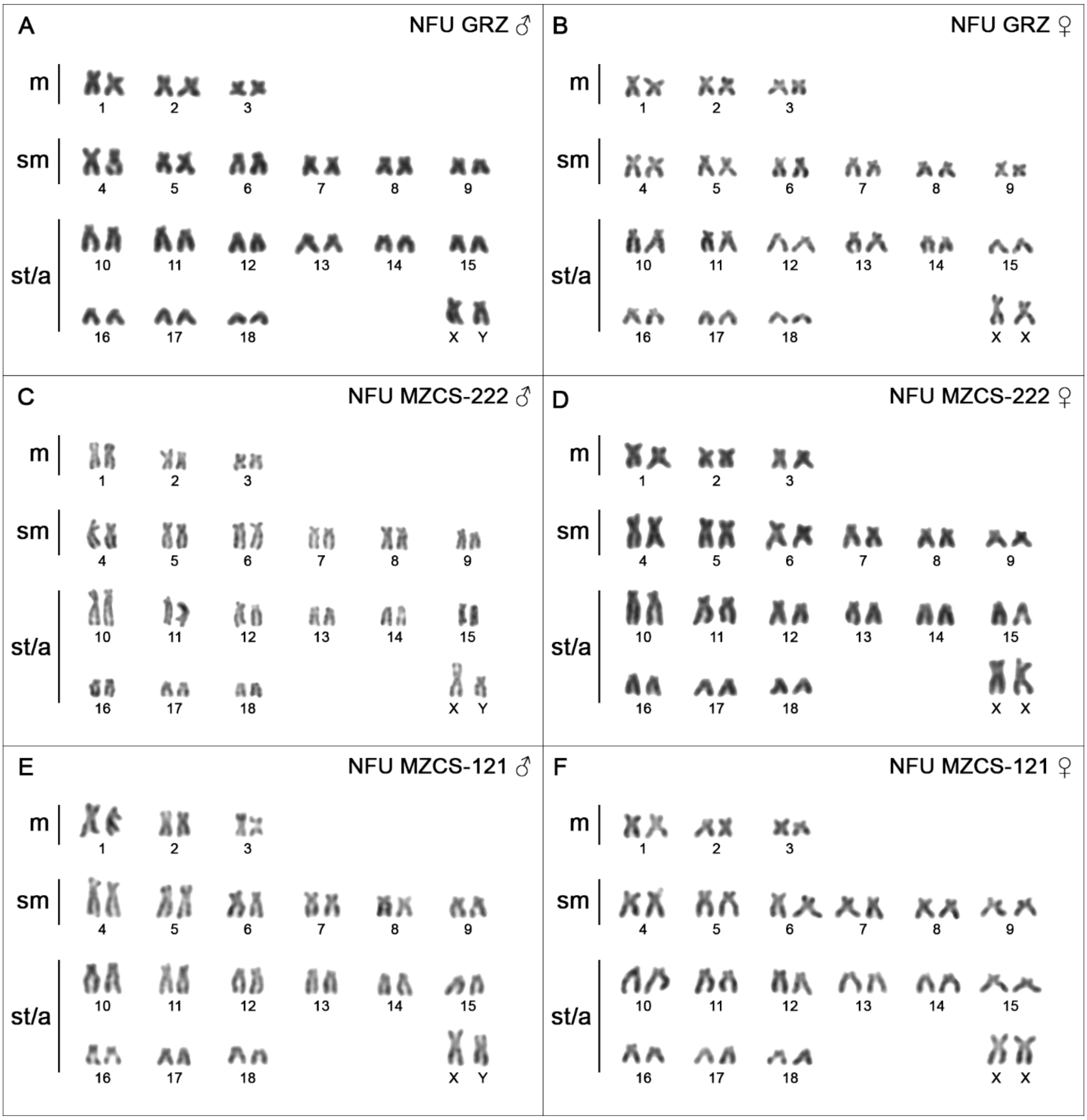


**S1 Fig. Karyotypes of *N. furzeri* males and females after Giemsa staining.** m = metacentric chromosome, sm = submetacentric chromosome, st/a = subtelocentric-to-acrocentric chromosome.

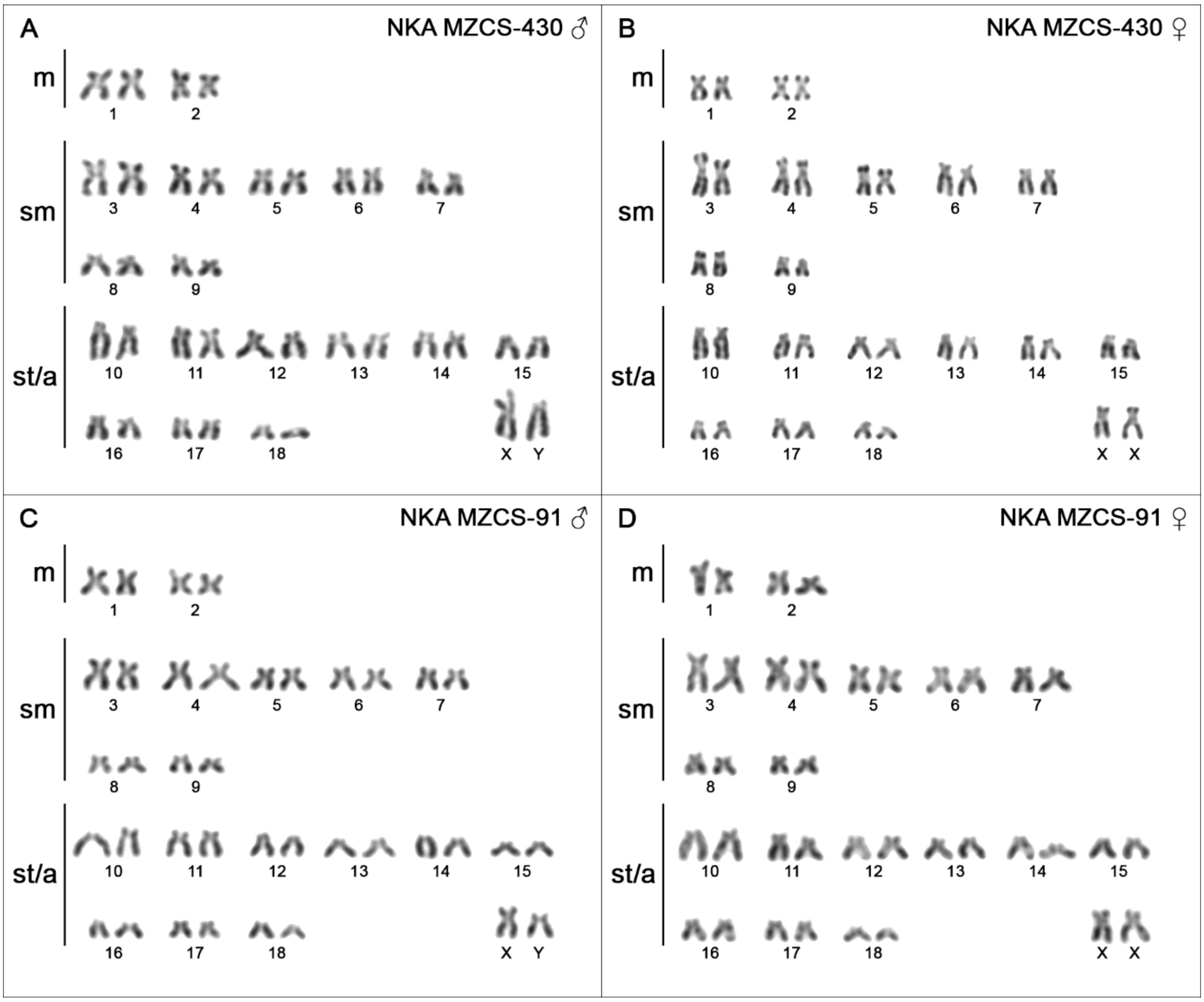


**S2 Fig. Karyotypes of *N. kadleci* males and females after Giemsa staining.** m = metacentric chromosome, sm = submetacentric chromosome, st/a = subtelocentric-to-acrocentric chromosome.

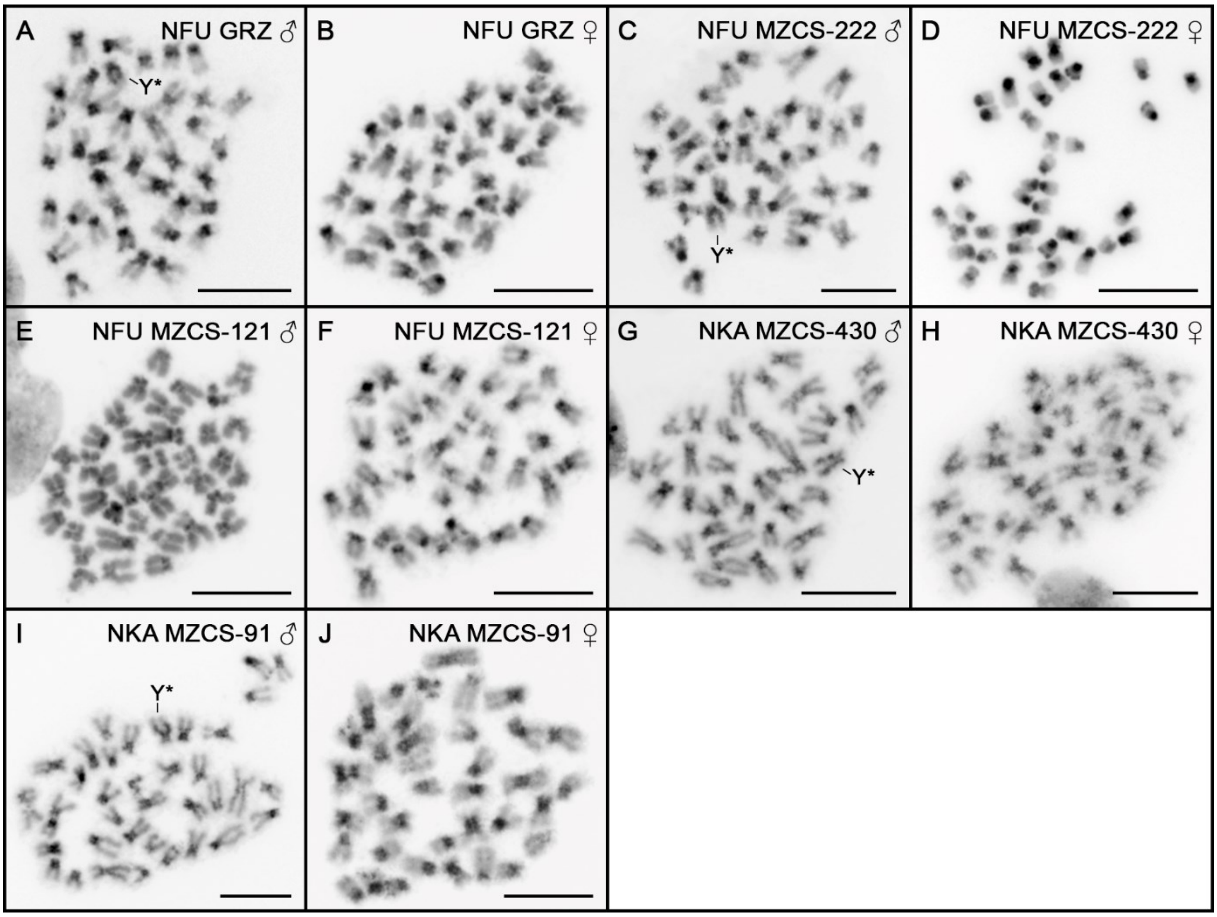


**S3 Fig. Mitotic metaphases of *N. furzeri* and *N. kadleci* after C-banding.** Chromosomes stained with DAPI (inverted picture). Putative Y sex chromosomes are marked. Scale bar = 10 µm.

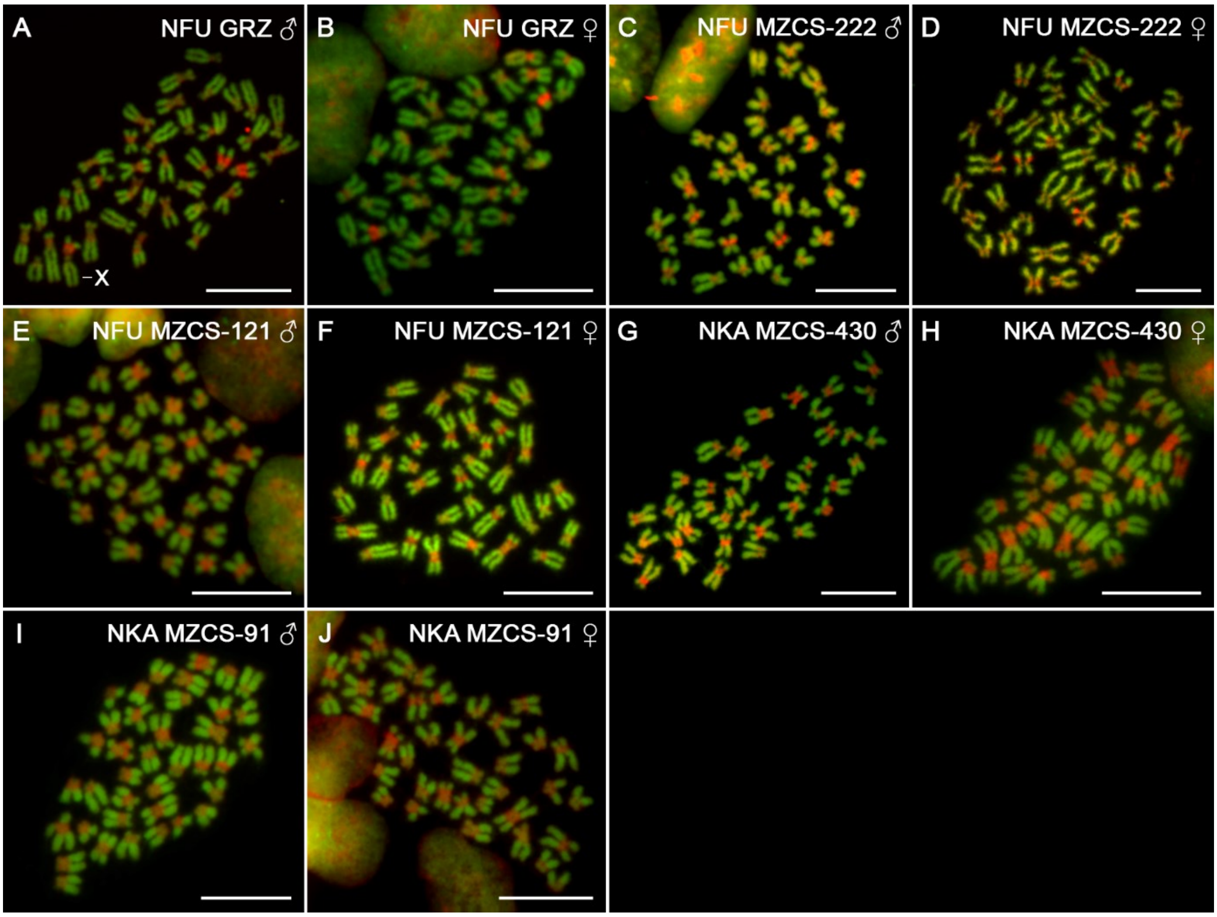


**S4 Fig. Mitotic metaphases of *N. furzeri* and *N. kadleci* after CMA3/DAPI staining.** For better contrast, images were pseudocoloured in red (for CMA3) and green (for DAPI). (**A**) X chromosome is marked based on morphology and strong CMA3-positive signal related to major rDNA site. Scale bar = 10 μm.

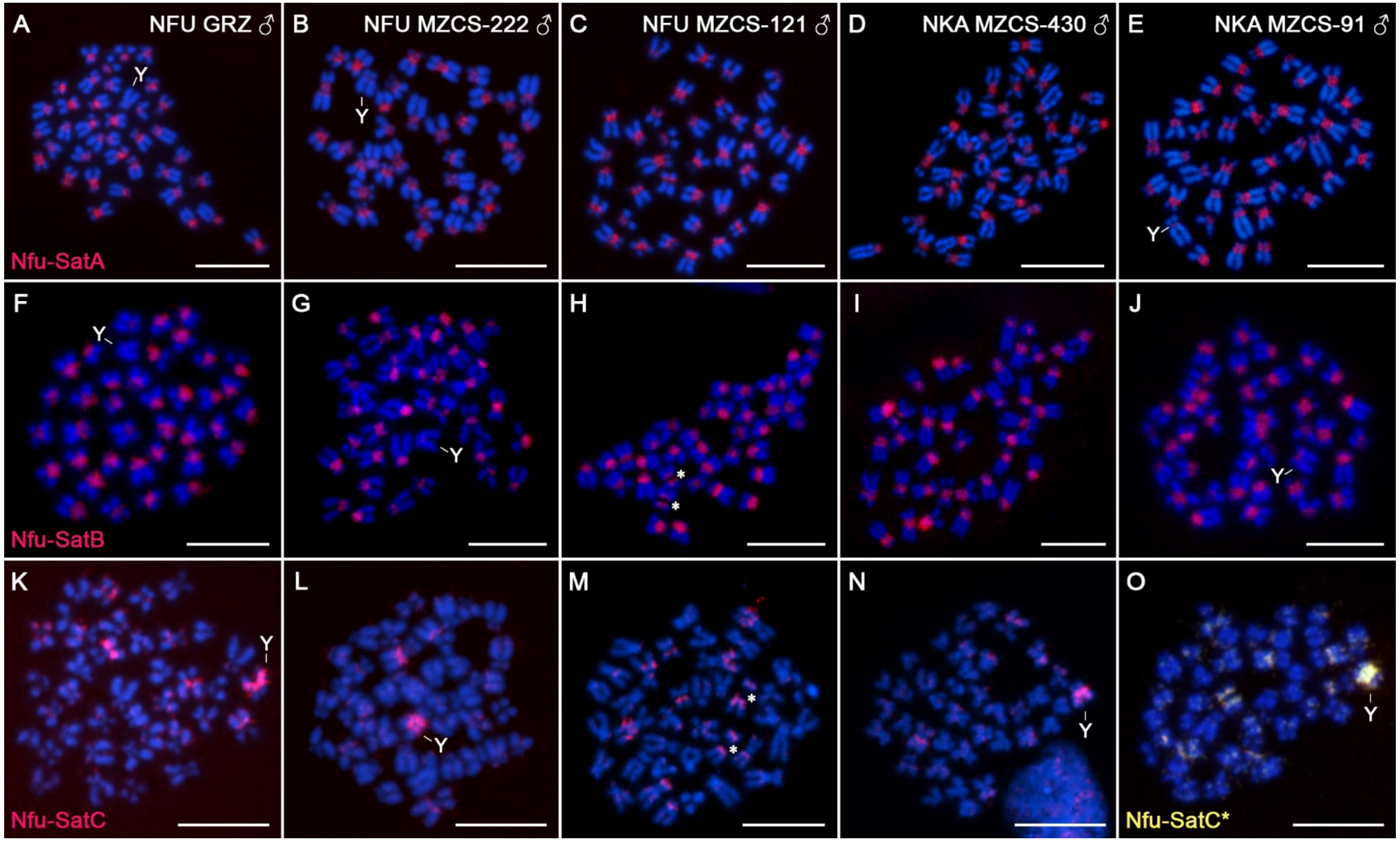


**S5 Fig. FISH with tandem repeats retrieved from RepeatExplorer analysis on male metaphases.** Red signals: (**A-E**) Nfu-SatA minisatellite, (**F-J**) Nfu-SatB satDNA, (**K-O**) Nfu-SatC satDNA. Yellow signals: FISH with two differently labeled Nfu-SatC probes derived from cloned fragments with different sequence compositions (sequences were deposited in GenBank under accession numbers OM542182 and OM542183); note that green and red signals are entirely overlapping, producing virtually uniform yellowish hybridization pattern. Y sex chromosomes are marked if possible. Asterisks point to population-specific gap in signal in one metacentric pair in NFU MZCS-121 (**H, M**). Chromosomes were counterstained with DAPI (blue). Scale bar = 10 µm.

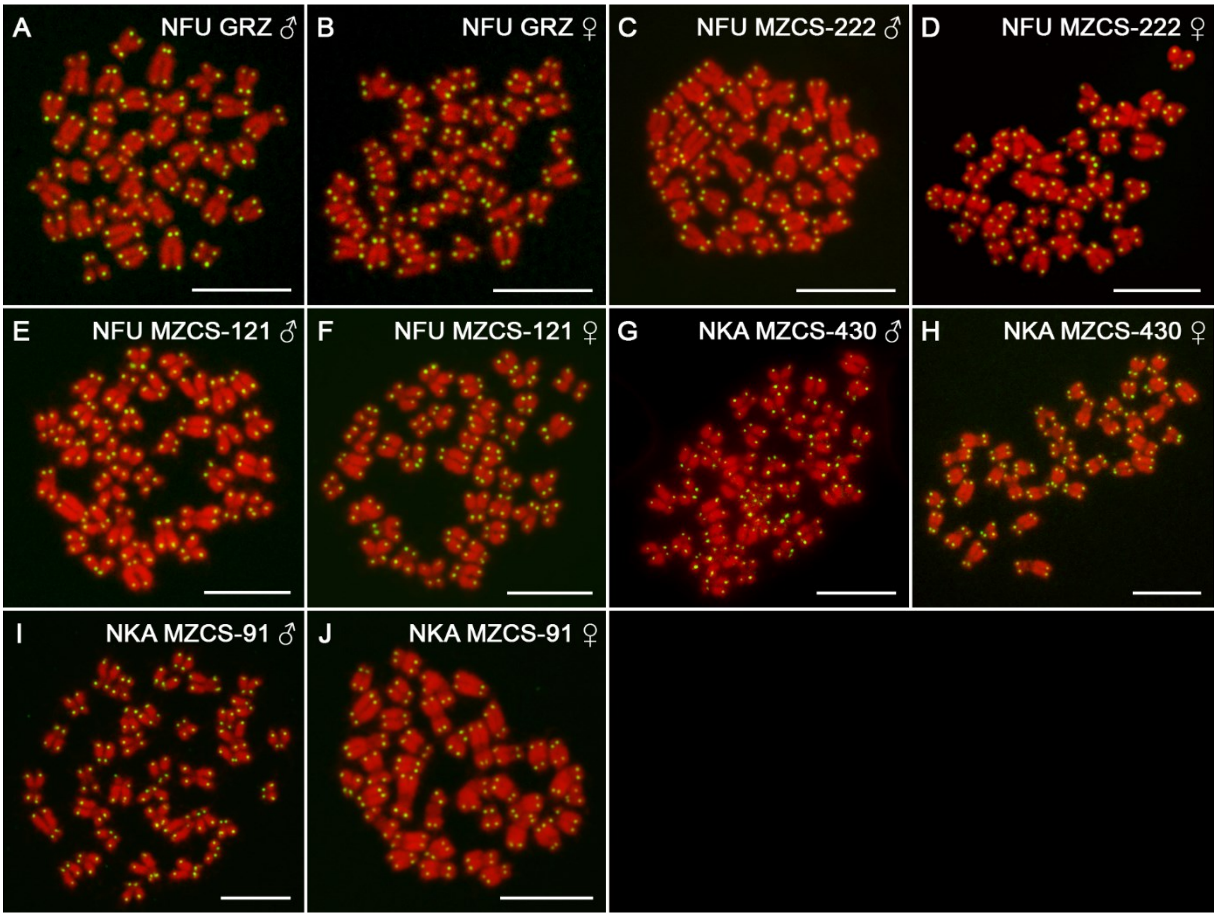


**S6 Fig. PNA-FISH with telomeric probe**. For better contrast, pictures were pseudocoloured in green (telomeric repeat probe) and red (DAPI). Scale bar = 10 µm.

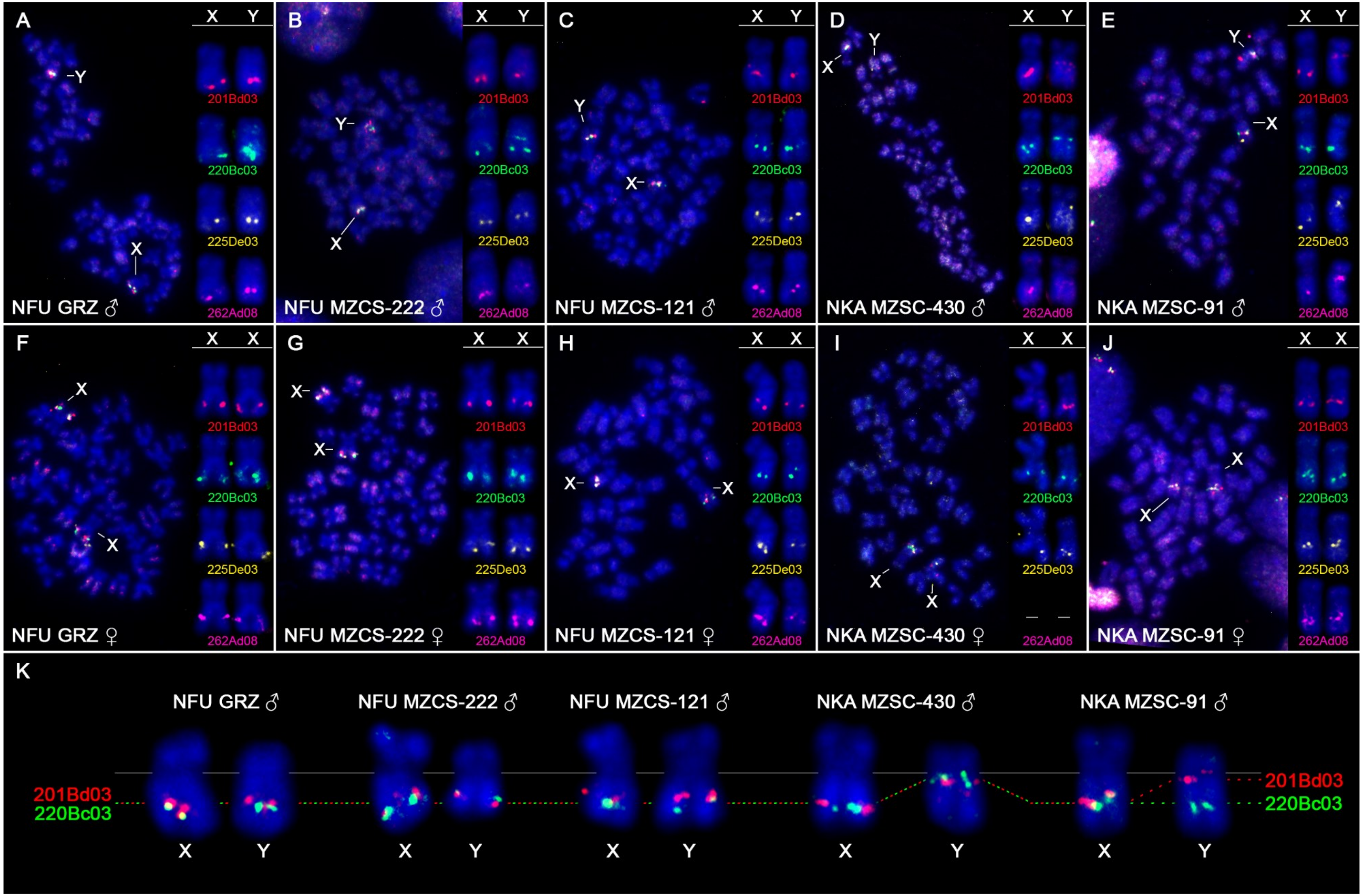
A
NFU GRZ ♂
B
NFU MZCS-222 ♂
C
NFU MZCS-121 ♂
D
NKA MZSC-430 ♂
E
NKA MZSC-91 ♂
F
NFU GRZ ♀
G
NFU MZCS-222 ♀
H
NFU MZCS-121 ♀
I
NKA MZSC-430 ♀
J
NKA MZSC-91 ♀
X
Y
201Bd03
220Bc03
225De03
262Ad08
K
NFU GRZ ♂
NFU MZCS-222 ♂
NFU MZCS-121 ♂
NKA MZSC-430 ♂
NKA MZSC-91 ♂
201Bd03
220Bc03

**S7 Fig. Physical map of selected XY-linked BAC clones.** Details on BAC clones are provided in S3 and S4 Tables. BAC bearing *gdf6* gene is 201Bd03. Note that in NKA MZCS-430 male (**D**), three out of four BACs less intense and diffuse signals on the Y chromosome, probably due to sequence divergence on this sex chromosome. In females of this population (**I**), we did not get reproducible signals of the 262Ad08 clone, however, its position is known from male X chromosome. Note the shift in position of all four BACs (201Bd03, 220Bc03, 225De03 and 262Ad08) on the Y chromosome of NKA MZCS-430 male, while only three BACs (201Bd03, 225De03 and 262Ad08) are shifted in NKA MZCS-91 male. (**K**) Detailed scheme showing this inversion polymorphism in *N. kadleci* narrowed to two informative BACs.

**S8 Fig. Sequential FISH experiments in *N. furzeri* males – full karyotypes.** The same metaphase plates were treated subsequently with different FISH protocols. i) BAC-FISH – dual-colour FISH with X-linked 220Bc03 BAC clone (green signal) and Y-linked 201Bd03 BAC (*gdf6y*-bearing clone; red signal); ii) single-colour FISH with Nfu-SatC probe (red signals); iii) dual-colour FISH with 5S (red signals) and 18S (green signals) rDNA probes. Note that 18S rDNA cluster is present on short arms of X chromosome but mostly not on Y chromosome. Some males from Limpopo population NFU MZCS-121 have exceptionally 18S rDNA site also on Y chromosome. iv) comparative genomic hybridization (CGH) performed sequentially after BAC-FISH on different set of metaphases – hybridization patterns of male (red) and female (green) genomic probes yielded uniform yellowish signal. No exclusive or biased accumulation of either probe was observed, failing thus to unmask the region of differentiation on Y chromosome. Note, however, differences between X and Y in the size of (peri)centromeric block of accumulated repetitive DNA. m = metacentric chromosome, sm = submetacentric chromosome, st = subtelocentric chromosome, st/a = subtelocentric-to-acrocentric chromosome.

**S9 Fig. Sequential FISH experiments in *N. furzeri* females – full karyotypes.** The same metaphase plates were treated subsequently with different FISH protocols. i) BAC-FISH – dual-colour FISH with X-linked 220Bc03 BAC clone (green signal) and Y-linked 201Bd03 BAC (*gdf6y*-bearing clone; red signal); ii) single-colour FISH with Nfu-SatC probe (red signals); iii) dual-colour FISH with 5S (red signals) and 18S (green signals) rDNA probes. Note that 18S rDNA cluster is present on short arms of X chromosomes but some females from Limpopo population NFU MZCS-121 lacked 18S rDNA site on one X homologue. m = metacentric chromosome, sm = submetacentric chromosome, st/a = subtelocentric-to-acrocentric chromosome.

**S10 Fig. Sequential FISH experiments in *N. kadleci* males and females – full karyotypes.** The same metaphase plates were treated subsequently with different FISH protocols. i) BAC-FISH – dual-colour FISH with X-linked 220Bc03 BAC clone (green signal) and Y-linked 201Bd03 BAC (*gdf6y*-bearing clone; red signal); ii) single-colour FISH with Nfu-SatC probe (red signals); iii) dual-colour FISH with 5S (red signals) and 18S (green signals) rDNA probes. Note that 18S rDNA cluster is present on short arms of X chromosome but not on Y chromosome. iv) comparative genomic hybridization (CGH) performed sequentially after BAC-FISH on different set of metaphases – hybridization patterns of male (red) and female (green) genomic probes yielded uniform yellowish signal. No exclusive or biased accumulation of either probe was observed, failing thus to unmask the region of differentiation on Y chromosome. Note, however, differences between X and Y in the size of (peri)centromeric block of accumulated repetitive DNA. m = metacentric chromosome, sm = submetacentric chromosome, st = subtelocentric chromosome, st/a = subtelocentric-to-acrocentric chromosome.

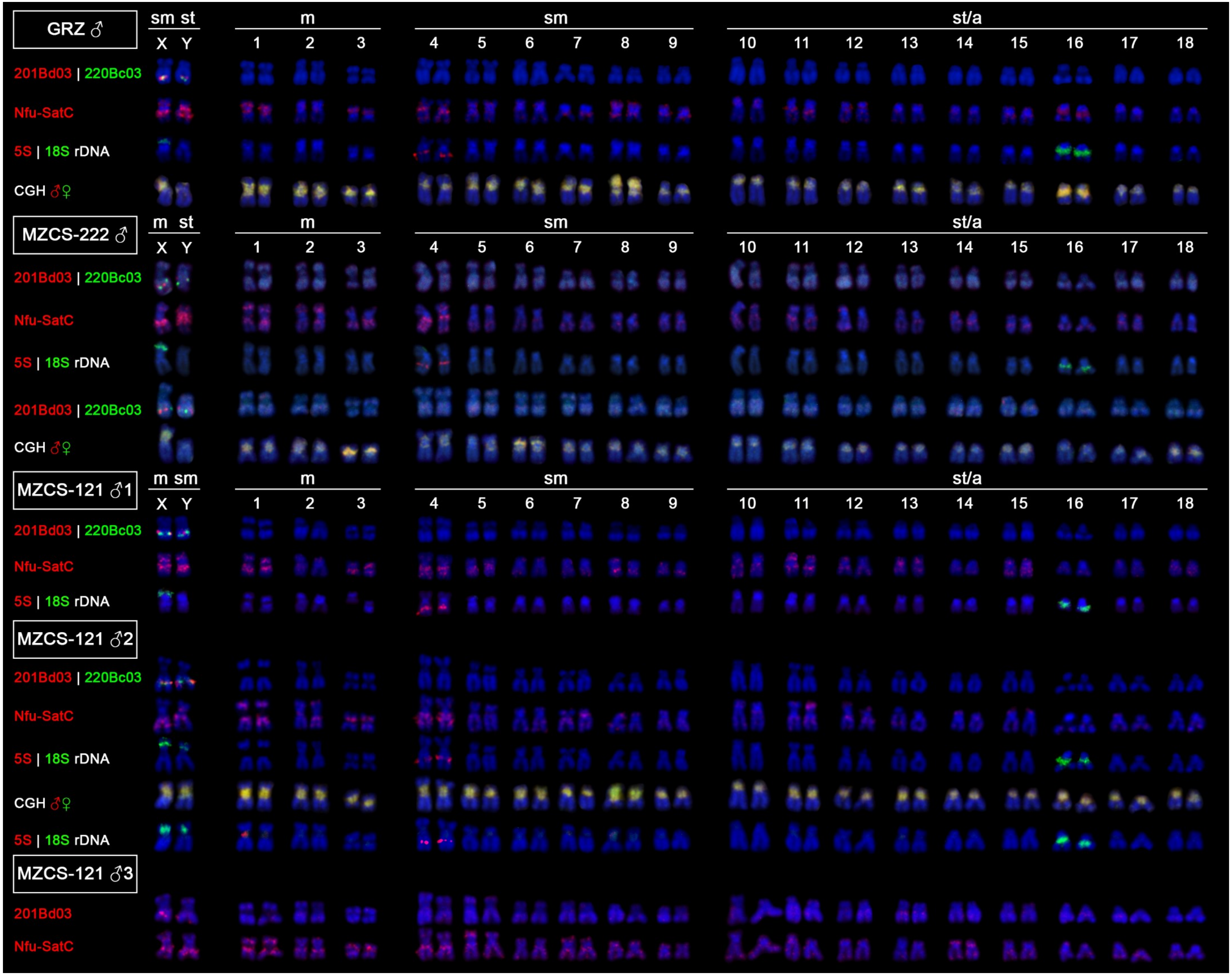

GRZ ♂
MZCS-222 ♂
MZCS-121 ♂1
MZCS-121 ♂2
MZCS-121 ♂3
sm st
m st
m sm
X Y
m
sm
st/a
1 2 3 4 5 6 7 8 9 10 11 12 13 14 15 16 17 18
201Bd03 | 220Bc03
Nfu-SatC
5S | 18S rDNA
CGH ♂♀
201Bd03

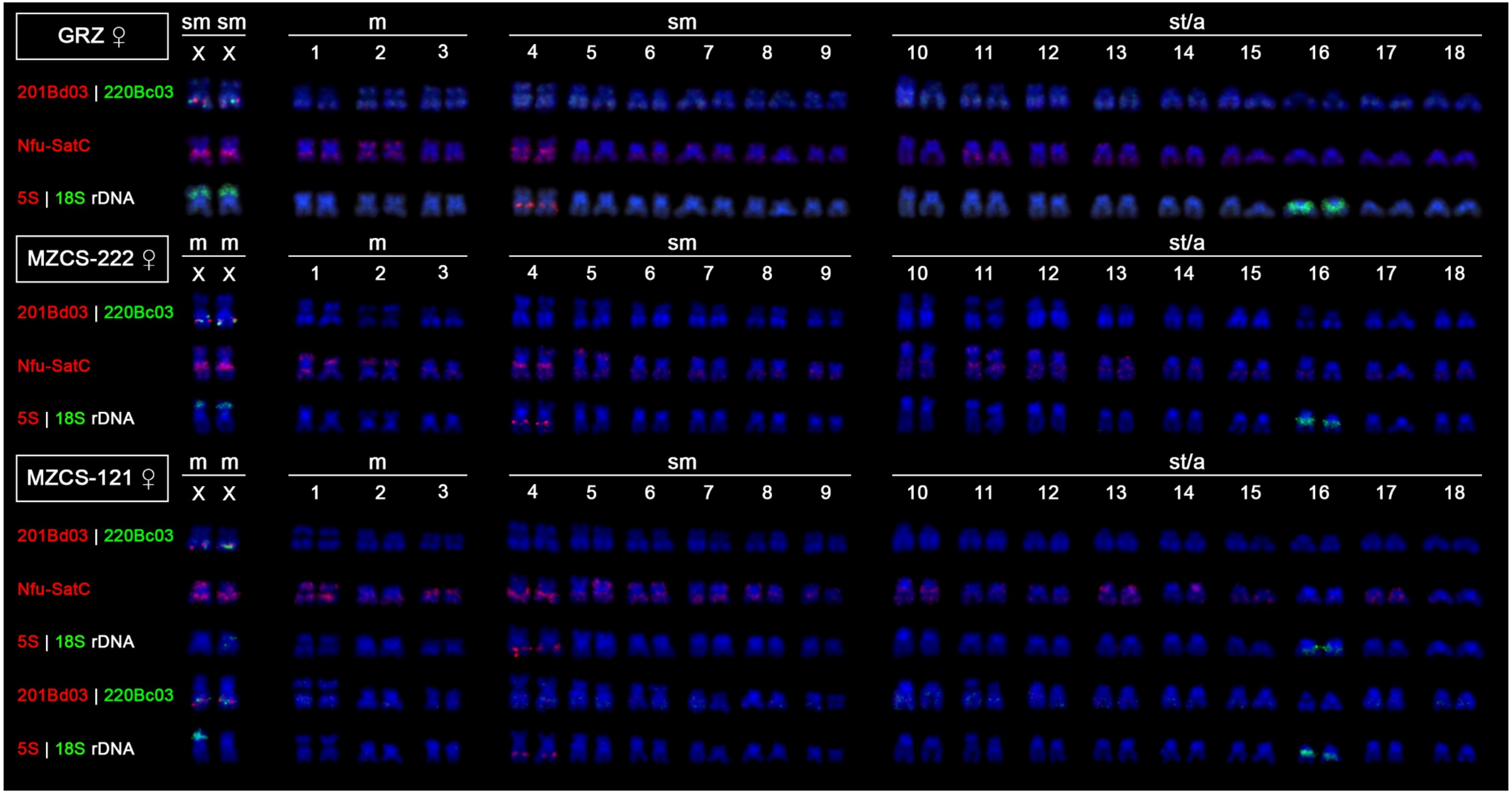
GRZ ♀
MZCS-222 ♀
MZCS-121 ♀
sm sm
m m
X X
m
sm
st/a
1 2 3 4 5 6 7 8 9 10 11 12 13 14 15 16 17 18
201Bd03 | 220Bc03
Nfu-SatC
5S | 18S rDNA

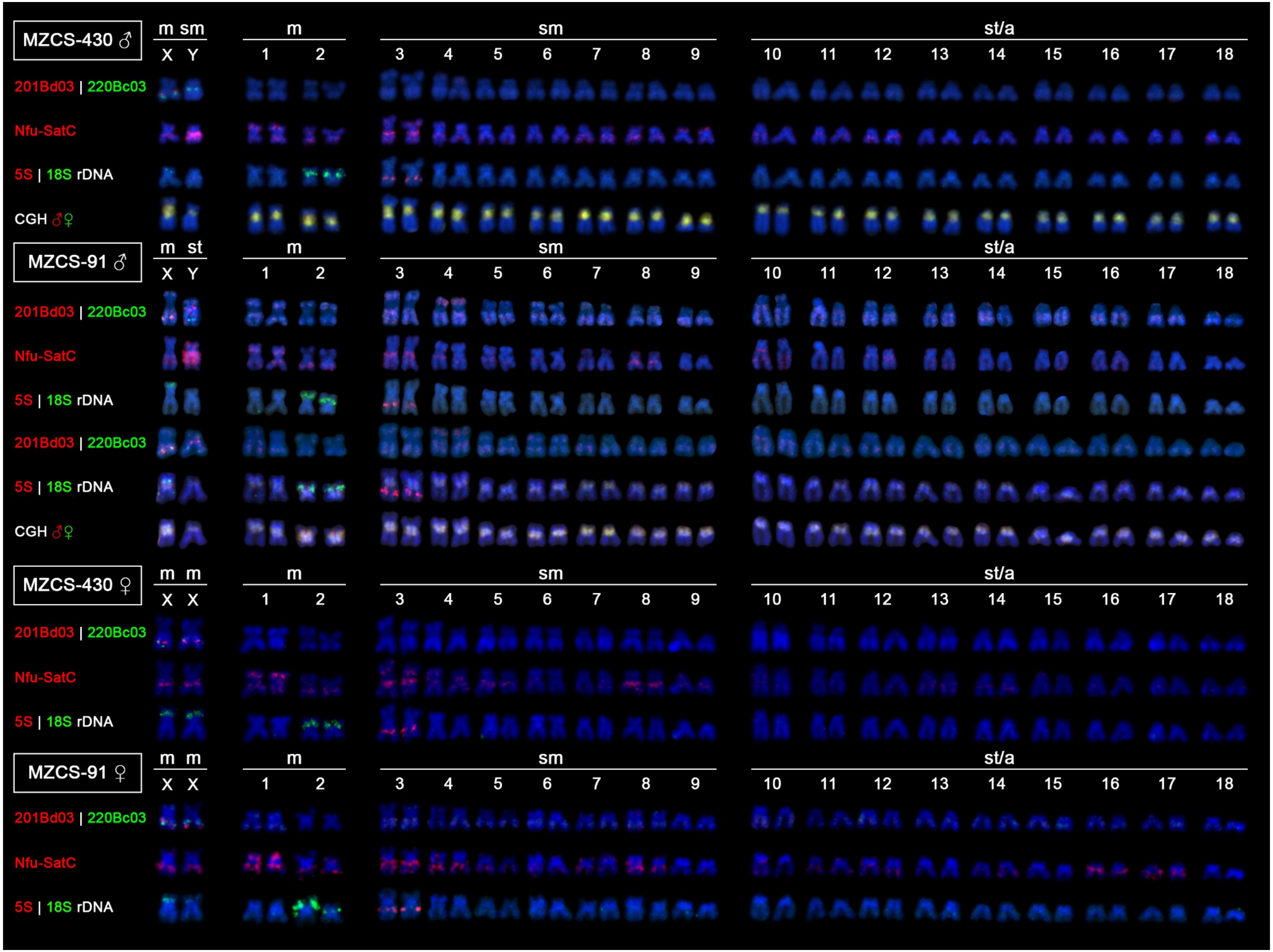
MZCS-430 ♂
m sm
X Y
m
1 2
sm
3 4 5 6 7 8 9
st/a
10 11 12 13 14 15 16 17 18
201Bd03 | 220Bc03
Nfu-SatC
5S | 18S rDNA
CGH ♂♀
MZCS-91 ♂
m st
X Y
m
1 2
sm
3 4 5 6 7 8 9
st/a
10 11 12 13 14 15 16 17 18
201Bd03 | 220Bc03
Nfu-SatC
5S | 18S rDNA
201Bd03 | 220Bc03
5S | 18S rDNA
CGH ♂♀
MZCS-430 ♀
m m
X X
m
1 2
sm
3 4 5 6 7 8 9
st/a
10 11 12 13 14 15 16 17 18
201Bd03 | 220Bc03
Nfu-SatC
5S | 18S rDNA
MZCS-91 ♀
m m
X X
m
1 2
sm
3 4 5 6 7 8 9
st/a
10 11 12 13 14 15 16 17 18
201Bd03 | 220Bc03
Nfu-SatC
5S | 18S rDNA

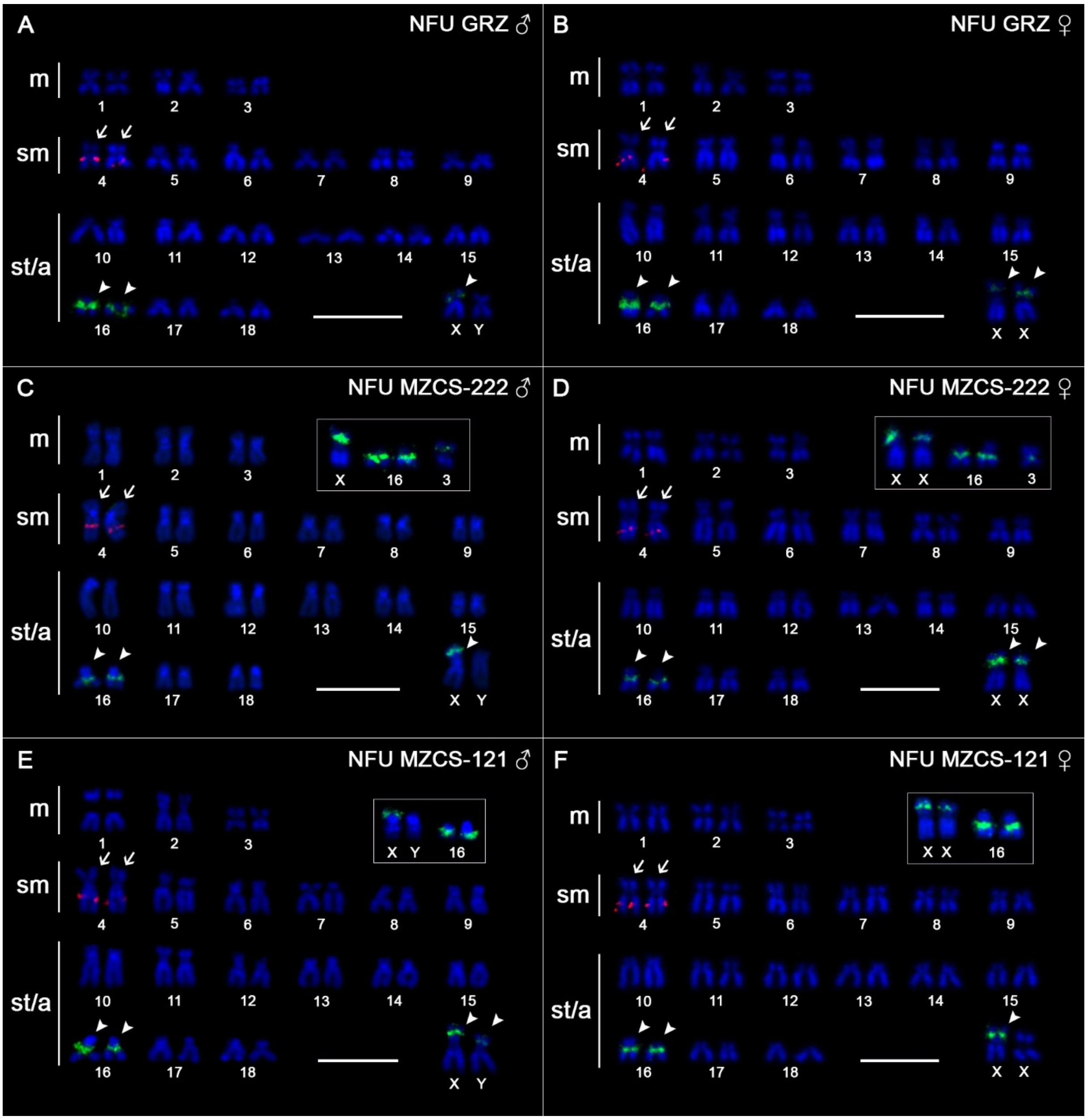


**S11 Fig. Karyotypes of *N. furzeri* after 5S/18S rDNA FISH.** 5S rDNA (red, arrows) and 18S rDNA (green, arrowheads); probes mapped on mitotic chromosomes. Chromosomes were counterstained with DAPI (blue). Note the presence of 18S rDNA site on X chromosomes and in NFU MZCS-121 males also on Y chromosome. Inter-individual variability in 18S rDNA sites is boxed. m = metacentric chromosome, sm = submetacentric chromosome, st/a = subtelocentric-to-acrocentric chromosome. Scale bar = 10 µm.

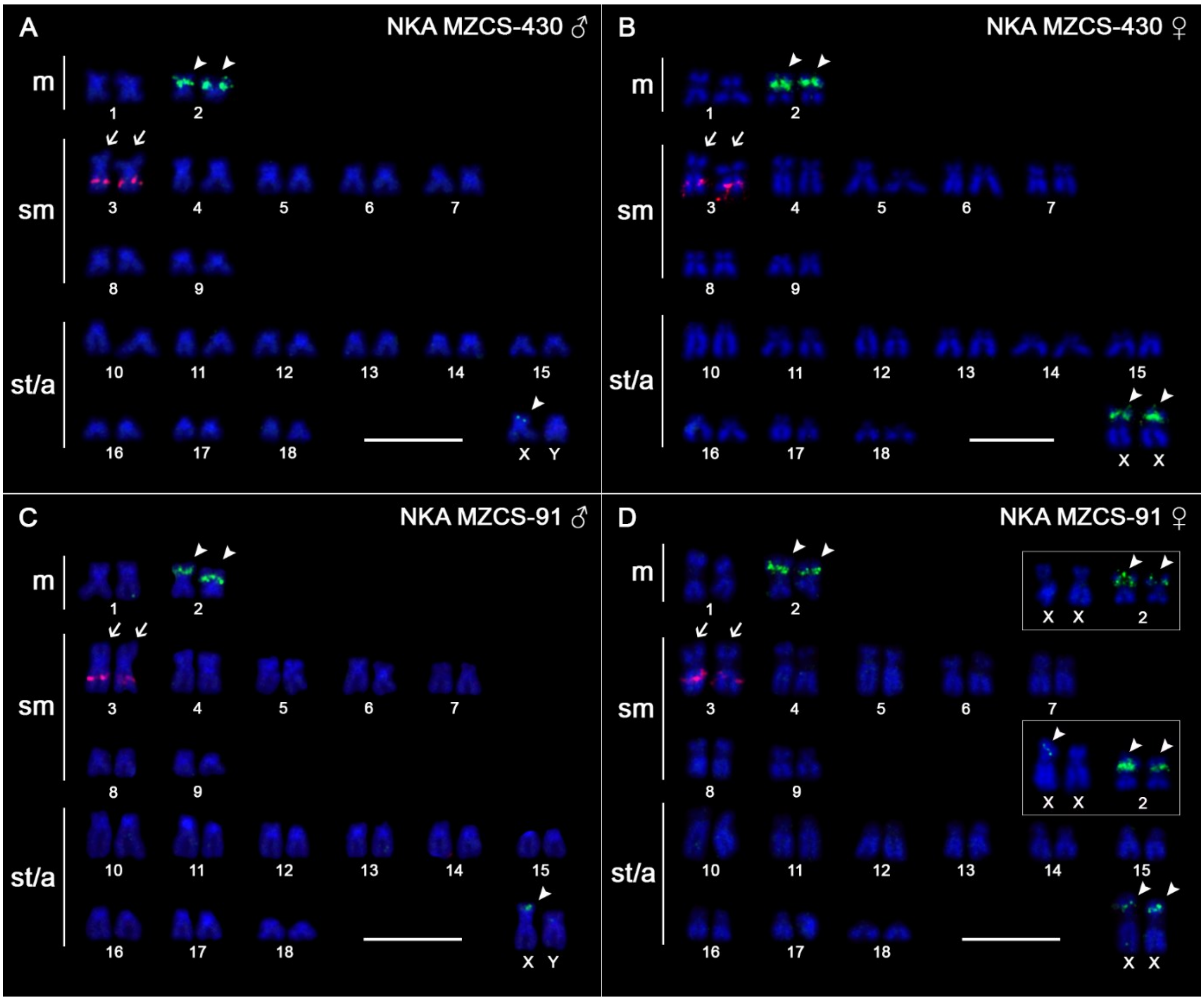


**S12 Fig. Karyotypes of *N. kadleci* after 5S/18S rDNA FISH.** 5S rDNA (red, arrows) and 18S rDNA (green, arrowheads); probes mapped on mitotic chromosomes. Chromosomes were counterstained with DAPI (blue). Note the presence of 18S rDNA site on X chromosomes. Inter-individual variability in 18S rDNA sites is boxed. m = metacentric chromosome, sm = submetacentric chromosome, st/a = subtelocentric-to-acrocentric chromosome. Scale bar = 10 μm.

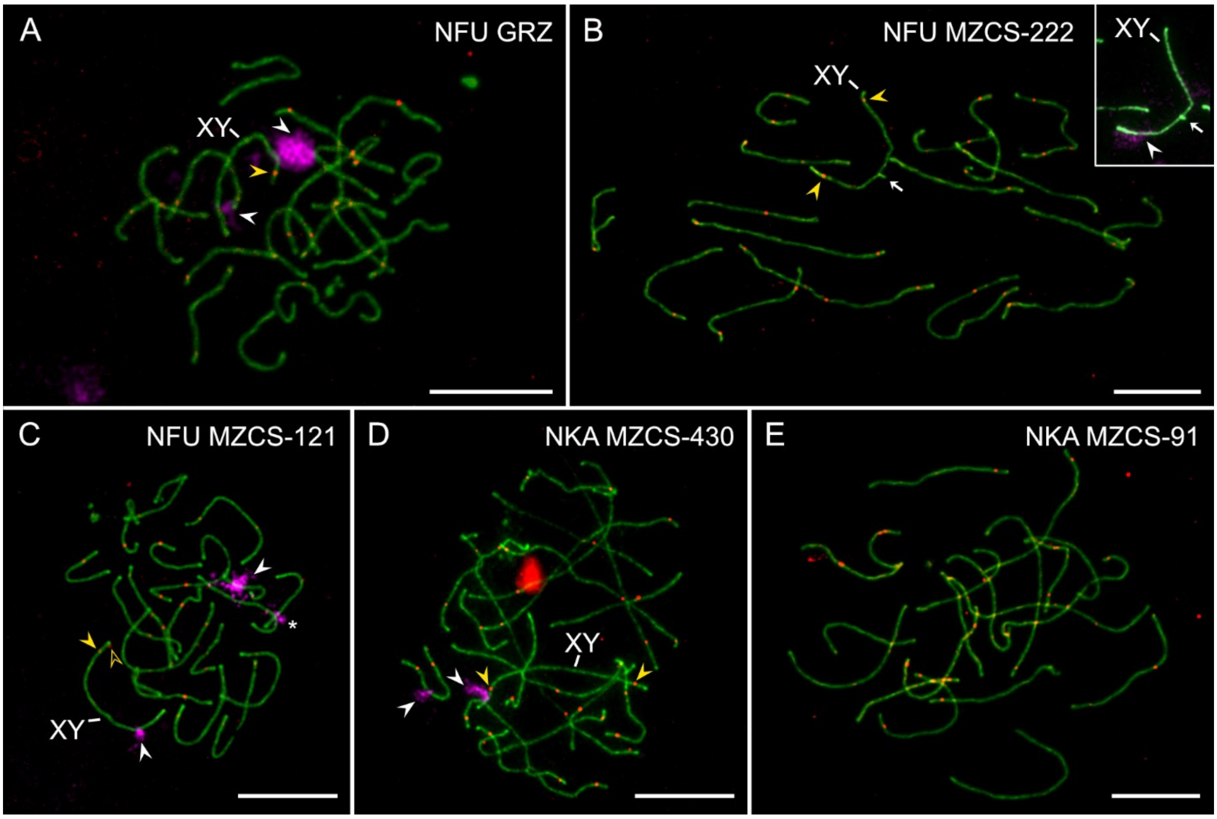


**S13 Fig. Male pachytene spreads after synaptonemal complex analysis and subsequent 18S rDNA FISH.** SCs were visualized by anti-SYCP3 antibodies (green), the recombination sites were identified by anti-MLH1 antibodies (red). Except for (**E**), slides were recycled for 18S rDNA FISH (magenta signals, white arrowheads). Based on patterns inferred from mitotic metaphases (see S8, S10-S12 Figs), XY bivalent is the larger from the two bivalents bearing 18S rDNA site. Yellow arrowheads point on MLH1 foci on XY sex bivalent. Empty yellow arrowheads denote putative MLH1 sites. Arrow (**B**) points on putative self-paired region found on XY bivalent based on 18S rDNA FISH confirmation (inset). Asterisk (**C**) points on highly probably false positive 18S rDNA signal (based on evidence on many meiotic and mitotic plates from individuals from NFU MZCS-121 population). Note that analysed males form 19 standard bivalents without asynapses, including XY sex bivalent, with a rare exception depicted in (**B**). Scale bar = 10 μm.

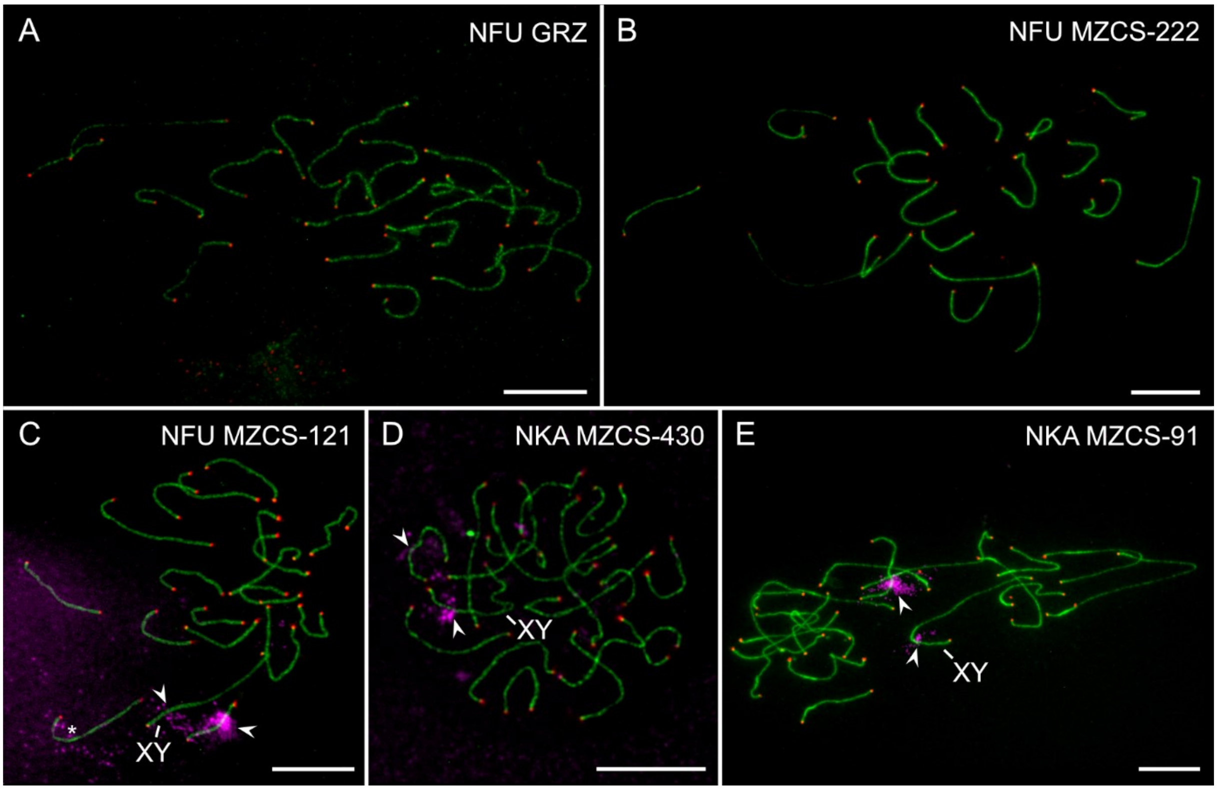


**S14 Fig. Male pachytene spreads after synaptonemal complex analysis and subsequent telomeric and 18S rDNA FISH.** SCs were visualized by anti-SYCP3 antibodies (green). SYCP3 immunostaining was followed by telomeric PNA FISH (red signals) (**A-E**) and in some cases additionally by 18S rDNA FISH (magenta signals, arrowheads) (**B**,**D**,**E**). Based on patterns inferred from mitotic metaphases (see S8, S10-S12 Figs), XY bivalent is the larger from the two bivalents bearing 18S rDNA site. Note that all telomeric signals are placed at the very ends of all bivalents which in the case of XY sex bivalent suggests synaptic adjustment in populations with heteromorphic sex chromosomes (i.e. all but NFU MZCS-121 where XY is often homomorphic). Scale bar = 10 µm.

### S1 Text. Characterization of major satellite DNA markers

Nfu-SatA represents a GC-rich minisatellite with a monomer size 77 bp which is the most abundant repetitive DNA in the genome of *N. kadleci* (approx. 9.6 %) and second abundant repeat class in the *N. furzeri* genome (3.5 %). For FISH, the probe covering the whole monomer length was generated by 5' labelling by Cy3 during synthesis (Generi Biotech, Hradec Králové, Czech Republic).

Monomer sequence of Nfu-SatA (5'-3'):

```
>Nothobranchius_furzeri_satellite_Nfu-SatA
GAGGAGCGTCTTGCCACATGTCAACACTGGGTCTGACTGAGCGCCTCCAGGAGAGAGGCCCA
AGCAAGACGCGCCCC
```

Nfu-SatB is a satDNA with 348 bp long motif which is the most abundant repeat found in the genome of *N. furzeri* (16 %) and second abundant repeat in *N. kadleci* (7.5 %). For FISH mapping, the probe was prepared from *N. kadleci* gDNA and the resulting approx. 1100 bp long fragment contained three tandemly arrayed monomers. DNA sequence of each monomer was identical with a consensus monomeric motif as identified by RepeatExplorer analysis in *N. furzeri* genome.

Consensual monomer sequence of Nfu-SatB (5'-3'):

```
>Nothobranchius_furzeri_satellite_Nfu-SatB
AGGTCCAATCAGGCTGAAACGTTACAGGAAGGTAGAATCTATTGAGTTGTAAGTGATCTGAA
CGTTTCATGATGATAGCACTTTCCTAAGGGGGTCAAACCATGTCCCAAAGGTCACCGGATCT
GGTGTCAATTTAAAGAAGTAGTCCAGTTACACATTTGTGGGCTTATAACTTGGCTTCTGAAT
GTCCTAGAGAGATCAGGTTTGTTTTAGTGTACTCCAATCAACAGCCCCTACAACCACAAAAA
GGAATGGAGTCATTAGCACTTATGGTTTGTAAATGGCACCCAGAATTATGTTGCACGAAGCA
CAATTTCACATCATTCTGGGTTCATTTGTGTGAAAACA
```

Nfu-SatC satDNA has a predicted monomer size 634 bp and encompasses 0.69 % of *N. furzeri* and 0.51 % of *N. kadleci* genomes. PCR amplification from *N. kadleci* genome yielded a set of fragments differing in size. After cloning and sequence analysis, we revealed high variability in fragment composition in terms of Nfu-SatC motif completeness and eventual presence of other sequences nested between incomplete Nfu-SatC monomers. In comparison to two previous repeats, Nfu-SatC hence presents considerably lower level of intragenomic sequence homogenization. For a FISH probe generation, we used fragments around 900 bp in size. Sequences from selected clones were deposited in GenBank under accession numbers OM542182 and OM542183.

Consensual monomer sequence of Nfu-SatC (5'-3'):

```
>Nothobranchius_furzeri_satellite_Nfu-SatC
TTTTTCACAGCTCCCTATTGGACTCCTTTTACCCAATCACCAGGAATCTCTGGTAAATAGCA
GCAGACAAGTTGAGGTCACTTGGTCTCCAGTACTGGCAGGTTTTGAAAAAAGGCGGGGCTTT
GGGAGCATGGCGAAATTCGCCATCACGCCATGGAAATACGTTTGCCTCTCATTTCTTCAGTT
```

ATCGTGATATCGCCACGAAATTTCTGGTGAGTGATCCTAGTCTGACCCCCAATAGAAATGGA
CAGCTGAAGTTTGTGGGCGTGGCCTATTTTCTGAAATAGCGCCCCCTAGGACCATTAAAACT
GTCAGCCCCAAGCCATGCTTTGACTGAGGATTACGAAATTTGGTACACTAATGTGGTGTCTC
AGGACCTACAAAAAAGTCTCTTGGAGCCAAGTGCAAAGTCGCACAGGAAGTCGGCCATTTTG
GTCCAAGTACGCGATTTAGTGGTTTTCGCACACGTTGTTTGGAGAGTGATGCTCCGTCGCCC
TTTTCACCAATCACCTTCAAACTTCTGCTATATACTCTTAAGACATAGAGGAAAAAATTCAA
CCGTCGGATTTTAAATAAGTATAAAGGTGTGGGCGTGGCTAAGCCTCAAACTTTGACCTTTC
GCCACGCCACTCTT

For Nfu-SatB and Nfu-SatC, we designed primers for their PCR amplification (S1 Table; for thermal profile, see S2 Table) and the probes for FISH were constructed from cloned fragments with verified sequence (for details, see Material and Methods section).

**S2 Text. Patterns of distribution of 18S rDNA**

Among individuals from *N. furzeri* populations (S11 Fig), the number of 18S rDNA clusters ranged from 3 to 5. A pair of signals was stably observed at pericentromeric position on st-a pair No. 16. Regarding sex chromosomes, 18S rDNA cluster was usually placed on the short arms of X chromosome while it lacked on Y chromosome. NFU GRZ population displayed stably three (males) and four (females) 18S rDNA signals (two autosomal and one-to-two X-linked ones), without any loci number polymorphism. In NFU MZCS-222 population, some individuals (6/15 males and 6/13 females) bore additional (fourth in males, fifth in females) signal on a single homologue from metacentric pair 3. Furthermore, in population NFU MZCS-121, intriguingly, eight out of 11 males possessed 18S rDNA signal of varied intensity also on Y sex chromosome (S11E Fig). Six out of nine females of this population, on the other hand, lacked 18S rDNA cluster on one X chromosome (S11F Fig). Probably with a contribution of this type of polymorphism together with variable copy number and spiralization of (peri)centromeric repeats, Xs sometimes appeared to slightly differ in size. In *N. kadleci*, the number of 18S rDNA clusters ranged from two to four. A single metacentric autosomal pair No. 2 invariably possessed 18S rDNA cluster on both homologues and remaining signals were, again, recorded on X chromosomes and never on Y chromosomes. Males and females from population NKA MZCS-430 and males from population NKA MZCS-91 lacked any loci number polymorphism, while females from NKA MZCS-91 population displayed three different patterns related to their X chromosomes. Out of nine females, five exhibited 18S rDNA loci on both Xs, three females displayed heterozygous condition (one homologue bore the signal) and a single female lacked any X-linked 18S rDNA cluster (S12D Fig). A summary of observed patterns is provided below.

| **Population** | ***N*** | **18S rDNA pattern** |
|---|---|---|
| NFU GRZ | 7 M | 7× 3 signals |
| | 7 F | 7× 4 signals |
| NFU MZCS-222 | 15 M | 9× 3 signals, 6× 4 signals |
| | 13 F | 7× 4 signals, 6× 5 signals |
| NFU MZCS-121 | 11 M | 3× 3 signals, 8× 4 signals |
| | 9 F | 6× 3 signals, 3× 4 signals |

| | | |
|---|---|---|
| NKA MZCS-430 | 6 M | 6× 3 signals |
| | 6 F | 6× 4 signals |
| NKA MZCS-91 | 7 M | 7× 3 signals |
| | 9 F | 5× 4 signals, 3× 3 signals, 1× 2 signals |

**S3 Text**. **General discussion about observed repetitive DNA patterns in *N. furzeri* and *N. kadleci* karyotypes.**

*The distribution of rDNA and telomeric repeats*

Telomeric FISH, which might be instrumental in revealing breakpoints of previous rearrangements, did not reveal any extra sites in addition to standard hybridization pattern (i.e. signals in all telomeres). Given the obvious high dynamics of *Nothobranchius* spp. karyotypes [Krysanov and Demidova 2018] it seems more likely that this pattern resulted from loss of telomeres during the mechanisms of rearrangements, or the site of rearrangement retained low copy number of telomeric sequence and/or the sequence was gradually eroded by mutations during evolution [Slijepcevic 1998, Ocalewicz 2013].

The number and location of 5S rDNA sites was found highly stable in our sampling. All individuals invariably possessed a single pair of submetacentric chromosomes bearing interstitial cluster of this multigene family. On the other hand, FISH with 18S rDNA probe for identification of major (45S rDNA) distribution revealed high level of both inter and intrapopulation variability, including size and presence/absence polymorphisms among homologues. These patterns could not result from the use of two different 18S rDNA probes in our experiments as they have virtually identical DNA sequence and multiple parallel experiments did not show any differences in obtained hybridization patterns. Most of the observed polymorphisms were related to sex chromosomes. A polymorphism unrelated to sex was represented by additional (fifth) 18S rDNA cluster in *N. furzeri* population MZCS-222 on a single homologue from metacentric pair No. 3, which is highly likely homeologous to 18S-bearing chromosome pair No. 2 in *N. kadleci* (where both homologues stably carry the signal). Notably, a stable 18S rDNA site present on st-a chromosome pair 16 in *N. furzeri* populations was absent from *N. kadleci* populations. It is therefore imaginable that we captured a process of rDNA site transition from one linkage group to another among closely related species and this might, to some extent, serve as cytogenetic marker for species genome identification in future studies. Mechanistically, these processes might be driven by ectopic (i.e. non-allelic) recombination resulting often from interaction of rDNA loci during interphase and/or from potential heterologous contacts during meiosis [Cazaux et al. 2011, Sember et al. 2015, Li et al. 2017]. Similar mechanisms might have contributed also to rDNA size heteromorphisms, where unequal crossing over between ectopic sites or between sister chromatids might drive repeat copy number variation [Charlesworth et al. 1994, Li et al. 2017]. Alternatively, rDNA cluster might move to chromosome 3 in NFU also by ectopic recombination. Finally, we cannot exclude the possibility that pair 16 in NFU could be homeologous to chromosome pair 2 in NKA as these chromosomes seem to be similar in size and pericentric inversion might have changed chromosome morphology in one species.

*Shared repeatome and patterns of tandem repeat distribution*

We sought to analyse genomes of both killifish species by RepeatExplorer2 pipeline, and we found that repeatomes of both species considerably overlap particularly regarding the most abundant tandemly repeated classes. We found that the most abundant repetitive DNA classes, Nfu-SatA and Nfu-SatB satellites are greatly amplified across almost all (peri)centromeres and thus overlap with blocks of constitutive heterochromatin revealed by C-banding. The only exception from the standard pattern found across our sampling is the Nfu-SatB distribution in NFU3 population where in both sexes a gap without hybridization signal was consistently found in centromere of one metacentric pair. This pattern suggests population-specific diversification regarding Nfu-SatB distribution.

### S4 Text. Reconstruction of X and Y cytogenetic maps in *N. furzeri* and *N. kadleci* populations

Cytogenetic maps of sex chromosomes were reconstructed based on measurements of: (i) absolute lengths of the X and Y for estimating the length difference between the sex chromosomes; (ii) the relative length and position of the markers of interest (i.e. BAC clones 201Bd03, 220Bc03; 18S rDNA; Nfu-SatC; repetitive elements revealed by CGH; p/q arm) on the X and Y chromosomes.

(i) Absolute lengths of the sex chromosomes were measured using the ImageJ software (https://imagej.nih.gov/ij/) (10 µm scale bar calibration). The % length difference between X and Y was calculated based on the mean values from six to ten chromosome pairs measured.

(ii) The measurements were made independently for each marker using the ImageJ software with the Levan plugin [Sakamoto and Zacaro 2009] which enables to analyse karyotype characteristics using relative length data. The tip of the short (p) arm of the sex chromosome was defined as the zero point. The chromosomal position and size of the marker region was measured and calculated relative to the total length of the chromosome. Mean values of relative lengths were estimated based on measurements from three to 13 chromosomes.

For details see the Dryad digital data repository (doi: 10.5061/dryad.4mw6m90ck).

| | NFU GRZ | | | | | | NFU MZCS-222 | | | | | | NFU MZCS-121 | | | | | | NFU MZCS-121 | | | | | |
|---|---|---|---|---|---|---|---|---|---|---|---|---|---|---|---|---|---|---|---|---|---|---|---|---|
| | **X** | | | **Y** | | | **X** | | | **Y** | | | **X** | | | **Y** | | | **X** | | | **Y** | | |
| | N | mean | SD | N | mean | SD | N | mean | SD | N | mean | SD | N | mean | SD | N | mean | SD | N | mean | SD | N | mean | SD |
| *p* arm | 13 | 34.28 | 1.83 | 13 | 26.34 | 1.85 | 10 | 43.86 | 3.66 | 7 | 24.97 | 1.50 | 7 | 46.73 | 1.32 | 7 | 47.10 | 2.31 | 8 | 40.79 | 1.93 | 8 | 37.10 | 1.89 |
| *q* arm | 13 | 65.72 | 1.83 | 13 | 73.66 | 1.84 | 10 | 56.14 | 3.66 | 7 | 75.03 | 1.50 | 7 | 53.27 | 1.32 | 7 | 52.91 | 2.31 | 8 | 59.21 | 1.93 | 8 | 62.90 | 1.89 |
| **201Bd03** | | | | | | | | | | | | | | | | | | | | | | | | |
| distance | 13 | 75.40 | 5.79 | 7 | 65.74 | 4.32 | 10 | 77.64 | 2.42 | 7 | 72.28 | 3.96 | 7 | 73.42 | 3.40 | 6 | 72.14 | 5.52 | 7 | 72.84 | 4.84 | 7 | 73.88 | 3.72 |
| **18S rDNA** | | | | | | | | | | | | | | | | | | | | | | | | |
| distance | 9 | 12.48 | 2.82 | – | – | – | 6 | 8.33 | 3.25 | – | – | – | 4 | 11.28 | 1.50 | – | – | – | 5 | 12.53 | 1.56 | 5 | 11.84 | 1.85 |
| length | 9 | 17.96 | 3.75 | – | – | – | 6 | 27.39 | 4.21 | – | – | – | 4 | 16.70 | 2.89 | – | – | – | 5 | 13.91 | 0.35 | 5 | 13.25 | 3.73 |
| **Nfu-SatC** (*p* arm) | | | | | | | | | | | | | | | | | | | | | | | | |
| distance | 2 | 18.16 | 4.52 | – | – | – | – | – | – | – | – | – | 7 | 21.75 | 2.39 | 7 | 12.49 | 1.79 | – | – | – | 5 | 19.58 | 3.34 |
| length | 2 | 13.58 | 2.54 | – | – | – | – | – | – | – | – | – | 7 | 19.51 | 1.75 | 7 | 17.28 | 4.16 | – | – | – | 5 | 15.59 | 1.20 |
| **Nfu-SatC** (*q* arm) | | | | | | | | | | | | | | | | | | | | | | | | |
| distance | 7 | 39.29 | 3.44 | 8 | 26.22 | 2.65 | 6 | 50.80 | 4.51 | 6 | 29.99 | 2.75 | 7 | 49.09 | 1.98 | 7 | 48.28 | 2.19 | 5 | 48.98 | 4.90 | 5 | 43.39 | 4.04 |
| length | 7 | 34.64 | 3.66 | 8 | 49.26 | 6.26 | 6 | 25.02 | 3.20 | 6 | 53.39 | 3.93 | 7 | 26.18 | 1.60 | 7 | 34.79 | 3.48 | 5 | 37.85 | 7.03 | 5 | 26.14 | 4.19 |
| **CGH repeats** | | | | | | | | | | | | | | | | | | | | | | | | |
| distance | 1 | 13.91 | – | – | – | – | 4 | 14.91 | 3.19 | – | – | – | – | – | – | – | – | – | 1 | 17.64 | – | 1 | 19.23 | – |
| length | 1 | 23.73 | – | – | – | – | 4 | 31.72 | 3.35 | – | – | – | – | – | – | – | – | – | 1 | 30.43 | – | 1 | 28.53 | – |

S4 Text – Table 1. Means of the relative length and position of four cytogenetic markers mapped on X and Y chromosomes from mitotic metaphases in *N. furzeri*. Mean values represent the % of the total length of the sex chromosome. Distance values: from the tip of the p-arm to the location of the marker; length values: from the tip to the end of the marker region. N – number of sex chromosomes measured; SD – standard deviation.

| | NKA MZCS-430 | | | | | | NKA MZCS-91 | | | | | |
|---|---|---|---|---|---|---|---|---|---|---|---|---|
| | **X** | | | **Y** | | | **X** | | | **Y** | | |
| | N | mean | SD | N | mean | SD | N | mean | SD | N | mean | SD |
| *p* arm | 12 | 43.42 | 2.19 | 7 | 26.29 | 2.58 | 13 | 45.67 | 2.04 | 11 | 24.74 | 2.05 |
| *q* arm | 12 | 56.58 | 2.19 | 7 | 73.71 | 2.58 | 13 | 54.33 | 2.04 | 11 | 75.26 | 2.05 |
| **201Bd03** | | | | | | | | | | | | |
| distance | 9 | 74.72 | 2.59 | 5 | 45.10 | 5.41 | 12 | 78.45 | 4.45 | 8 | 34.78 | 5.83 |
| **220Bc03** | | | | | | | | | | | | |
| distance | – | – | – | – | – | – | – | – | – | 4 | 73.68 | 1.52 |
| **18S rDNA** | | | | | | | | | | | | |
| distance | 9 | 12.81 | 3.54 | – | – | – | 7 | 12.00 | 3.30 | – | – | – |
| length | 9 | 16.20 | 2.34 | – | – | – | 7 | 12.71 | 1.97 | – | – | – |
| **Nfu-SatC** (*p* arm) | | | | | | | | | | | | |
| distance | – | – | – | – | – | – | – | – | – | 4 | 9.12 | 3.07 |
| length | – | – | – | – | – | – | – | – | – | 4 | 11.27 | 2.05 |
| **Nfu-SatC** (*q* arm) | | | | | | | | | | | | |
| distance | 6 | 55.40 | 2.94 | 4 | 26.73 | 3.39 | 5 | 51.28 | 3.44 | 5 | 29.37 | 4.46 |
| length | 6 | 14.79 | 1.35 | 4 | 48.58 | 6.91 | 5 | 32.66 | 4.69 | 5 | 50.71 | 4.10 |
| **CGH repeats** | | | | | | | | | | | | |
| distance | 3 | 21.76 | 1.30 | – | – | – | 6 | 19.34 | 2.74 | – | – | – |
| length | 3 | 31.31 | 2.42 | – | – | – | 6 | 36.35 | 1.91 | – | – | – |

S4 Text – Table 2. Means of the relative length and position of five cytogenetic markers mapped on X and Y chromosomes from mitotic metaphases in *N. kadleci*. Mean values represent the % of the total length of the sex chromosome. Distance values: from the tip of the p-arm to the location of the marker; length values: from the tip to the end of the marker region. N – number of sex chromosomes measured; SD – standard deviation.

| | NFU GRZ | | | NFU MZCS-222 | | | NFU MZCS-121 | | | NFU MZCS-121 | | | NKA MZCS-430 | | | NKA MZCS-91 | | |
|---|---|---|---|---|---|---|---|---|---|---|---|---|---|---|---|---|---|---|
| | N | mean | SD | N | mean | SD | N | mean | SD | N | mean | SD | N | mean | SD | N | mean | SD |
| **X** | 7 | 4.02 | 0.63 | 6 | 4.03 | 0.66 | 7 | 4.66 | 0.52 | 7 | 4.69 | 0.75 | 6 | 4.28 | 0.49 | 10 | 4.33 | 0.49 |
| **Y** | 7 | 3.23 | 0.30 | 6 | 2.90 | 0.38 | 7 | 4.22 | 0.48 | 7 | 4.21 | 0.75 | 6 | 3.89 | 0.43 | 10 | 3.82 | 0.43 |

S4 File – Table 3. Means of the absolute lengths of sex chromosomes in µm. N – number of sex chromosomes measured; SD – standard deviation.

### S1 Table. Primers used for repetitive DNA amplification in this study.

| Target | Forward primer (5 '-3 ') | Reverse primer (5 '-3 ') | Reference |
|---|---|---|---|
| *5S rDNA* | TACGCCCGATCTCGTCCGATC | CAGGCTGGTATGGCCGTAAGC | [Martins and Galetti 1999] |
| *18S rDNA* | CCGAGGACCTCACTAAACCA | CCGCTTTGGTGACTCTTGAT | [Cioffi et al. 2009] |
| Nfu-SatB | TTTCCTAAGGGGGTCAAACC | CTGGGTGCCATTTACAAACC | this study |
| Nfu-SatC | TGACCCCCAATAGAAATGGA | CGTGTGCGAAAACCACTAAA | this study |

### S2 Table. PCR thermal profiles used in this study.

| PCR step | 5S rDNA | 18S rDNA | Nfu-SatB, Nfu-SatC |
|---|---|---|---|
| Initial denaturation | 94 °C / 3 min | 95 °C / 5 min | 95 °C / 4 min |
| a. Denaturation | 94 °C / 30 s | 95 °C / 1 min | 95 °C / 1 min |
| b. Primer annealing | 60 °C / 1 min | 60 °C / 1 min | 60 °C / 1 min |
| c. Extension | 72 °C / 2 min | 72 °C / 1 min 30 s | 72 °C / 1 min 30 s |
| d. Final extension | 10 min | 10 min | 10 min |
| Number of cycles ( a.-c.) | 35× | 35× | 35× |
| Reference | [Martins and Galetti 1999] | [Yano et al. 2017] | Based on [Yano et al. 2017] |

### S3 Table. Selected BAC clones (following information from Reichwald et al. 2015, supplement S1I)

| BAC ID | ENA Accession | Position in XY assembly | | Length (bp) | Sex chromosome |
|---|---|---|---|---|---|
| | | Start | End | | |
| 262Ad08 | LN877285 | 37 520 | 157 492 | 119 972 | Y |
| 225De03 | LN877277 | 54 897 | 194 528 | 139 631 | X |
| 201Bd03 (*gdf6*) | LN877265<br>LN877266 | 142 614 | 277 904 | 135 290 | X |
| 220Bc03 | LN877275 | 150 456 | 275 228 | 124 772 | X |

### S4 Table. Primers for BAC clone verification

| BAC clone ID | Forward primer | Reverse primer |
|---|---|---|
| 262Ad08 | GCAGAAATCATCCTCACCACCA | GGTTGCTGTTGTGTCCTTGG |
| 225De03 | AATCATCCTTGCGGCTCTGT | GGTGGTGCTCTTTTCTGGGT |
| 201Bd03 (*gdf6*) | TTACGCGCCTTGGTTTTGAA | GTGGATCAATCGTTCAGCCC |
| 220Bc03 | GGCAGCGTCTTTGTTCACAT | ATCTCTGCTCTCCCCTCCAG |
